# GoldRush: A *de novo* long read genome assembler with linear time complexity

**DOI:** 10.1101/2022.10.25.513734

**Authors:** Johnathan Wong, Lauren Coombe, Vladimir Nikolić, Emily Zhang, Ka Ming Nip, Puneet Sidhu, René L Warren, Inanç Birol

## Abstract

**Motivation:** Current state-of-the-art long read *de novo* genome assemblers follow the Overlap Layout Consensus (OLC) paradigm, an O(n^2^) algorithm in its naïve implementation. While the most time- and memory-intensive step of OLC —the all-vs-all sequencing read alignment process— was improved and reimplemented in modern long read assemblers, these tools still often require excessive computational memory when assembling a typical 50X human genome dataset.

**Results:** Here we present GoldRush, a *de novo* genome assembly algorithm with linear time complexity in the number of input long sequencing reads. We tested GoldRush on Oxford Nanopore Technologies datasets with different base error profiles describing the genomes of three human cell lines (NA24385, HG01243 and HG02055), *Oryza sativa* (rice), and *Solanum lycopersicum* (tomato). GoldRush achieved NGA50 lengths of 18.3-22.2 Mbp for the three human datasets, with two of the three assemblies having the fewest extensive misassemblies, and NGA50 lengths of 0.3 and 2.6 Mbp for the 373 Mbp and 824 Mbp genomes of rice and tomato, respectively. Further, GoldRush assembled all genomes within a day, using at most 54.5 GB of RAM. These results demonstrate that our algorithm and new assembly paradigm can be used to assemble large genomes *de novo* efficiently in compute memory space, with resulting assembly contiguity comparable to that of state-of-the-art OLC genome assemblers.

**Availability:** https://github.com/bcgsc/goldrush

## 1 Introduction

Short-read genome assembly methods typically struggle to resolve sequence repeats and often fail to generate assemblies that reach chromosome-scale (Treangen & Salzberg, 2012). Both prokaryotic and eukaryotic genomes can contain a large proportion of repeats (Haubold & Wiehe, 2006), with the human genome estimated to be 66-69% repetitive (de Koning et al., 2011). Thus, it is imperative that these repetitive regions be sufficiently resolved for a successful *de novo* assembly. Innovations in bioinformatics have emerged to address this challenge, leveraging long-range evidence afforded by various data types, including linked reads (Afshinfard et al., 2022; Coombe et al., 2018), Hi-C (Putnam et al., 2016), and long sequencing reads (Coombe et al., 2021; Qin et al., 2019).

Long read sequencing technology has become increasingly prevalent in recent years. Sequencing throughput, affordability and the long read lengths are some of the key reasons (Adewale, 2020). The long read lengths, ranging from kilobases to megabases, enable better resolution of structural variants (Sakamoto et al., 2021) and long repeats (Bongartz, 2019). Long reads also enable correct and accurate identification of tandem repeat expansions(Chiu et al., 2021).

Oxford Nanopore Technologies (ONT), Plc. (Oxford, UK) and Pacific Biosciences (PacBio), Inc. (Menlo Park, USA) are currently the two preeminent providers of commercial long read sequencing technology. PacBio generally produces long reads with lower base errors (<1% for HiFi), but with shorter read lengths (typically averaging 10-25 kbp) (Hon et al., 2020) compared to that of ONT (typically 10-100+ kbp) (Dohm et al., 2020). Yet, high error rates (1-13%) (*New Research Algorithms Yield Accuracy Gains for Nanopore Sequencing*, 2020; *Q20+ Chemistry for Single Molecule Accuracy of 99% and Higher*, n.d.) in ONT reads remain a challenging obstacle to *de novo* assembly. Unlike the *de novo* assembly strategies designed for short reads, both long read sequencing technologies – and especially ONT, require algorithms and data structures that can accommodate mismatches and indels in the sequencing data.

Most long read assemblers follow the Overlap Layout Consensus paradigm (OLC), a quadratic run time algorithm in its naïve implementation. OLC consists of three steps. The first step, overlap, typically generates an overlap graph by computing the pairwise alignment of all reads. As datasets often contain tens of millions of reads, finding and storing the detected overlaps is the most computationally- and memory-intensive step in the OLC paradigm, and has been the target of recent innovative algorithms (Kolmogorov et al., 2019; Ruan & Li, 2020; Shafin et al., 2020). In the second step, layout, the generated read overlap graph is traversed to produce contigs, or contiguous sequences, that reconstruct the underlying genome. The last step, consensus, uses read alignments to infer the most likely nucleotide bases across contigs, and corrects the sequences accordingly (Z. Li et al., 2012; Wajid & Serpedin, 2012).

In recent years, a number of OLC-based *de novo* long read assemblers have been developed that leverage the long-range evidence provided by the technology. These tools include Flye (Kolmogorov et al., 2019), Redbean (Ruan & Li, 2020), and Shasta (Shafin et al., 2020). Each tool brings a different innovation to the table, with implementations of the OLC paradigm aiming to reduce the computational cost and address the high error rates of long reads. For instance, Flye clusters the long reads that are likely to originate from the same genomic locus in a preprocessing step to reduce the number of pairwise comparisons (Kolmogorov et al., 2019). Redbean segments each read into 256 bp tiling subsequences, reducing the dynamic programming matrix to a size of 65536 (=256×256) and speeding up the pairwise alignment process (Ruan & Li, 2020). On the other hand, to address the high error rate of long reads, Shasta compresses all homopolymers in the reads using run-length encoding, thereby removing all homopolymer expansion errors, one of the more common error types in the ONT data, and improving the accuracy of alignments in the overlap step of OLC (Shafin et al., 2020). While these optimizations have reduced the time it takes to assemble the long sequencing reads and ultimately improve upon the quality of the generated genome assemblies, these tools have a large memory footprint, requiring upwards of several hundred gigabytes of RAM for assembling a typical 50X human genome dataset.

Here we present GoldRush, a memory-efficient long read assembler that runs in linear time in the number of reads. GoldRush is implemented as a modular pipeline with four main steps. We show that GoldRush produces contiguous and correct assemblies with a low memory footprint, and does so without read-to-read alignments, marking an important paradigm shift in the assembly of long sequencing reads.

## 2 Methods

The GoldRush assembly pipeline consists of four steps: GoldRush-Path, GoldRush-Edit, Tigmint-long (Coombe et al., 2021; Jackman et al., 2018), and GoldRush-Link (Coombe et al., 2021). GoldRush-Path (GR-Path) first iterates through the long reads and generates a “golden path”, selected reads with a ~1X representation of the genome of interest. Because the output of GR-Path is a set of raw reads, the golden path is first polished using GoldRush-Edit (GR-Edit) to correct base errors. Next, misassemblies (due to chimeric reads) are corrected using Tigmint-long (Coombe et al., 2021; Jackman et al., 2018). Finally, the corrected golden path is scaffolded using GoldRush-Link (GR-Link) to produce the output assembly (Coombe et al., 2021) (Fig. 1a and Supplementary Fig. S1).

**Fig 1.**
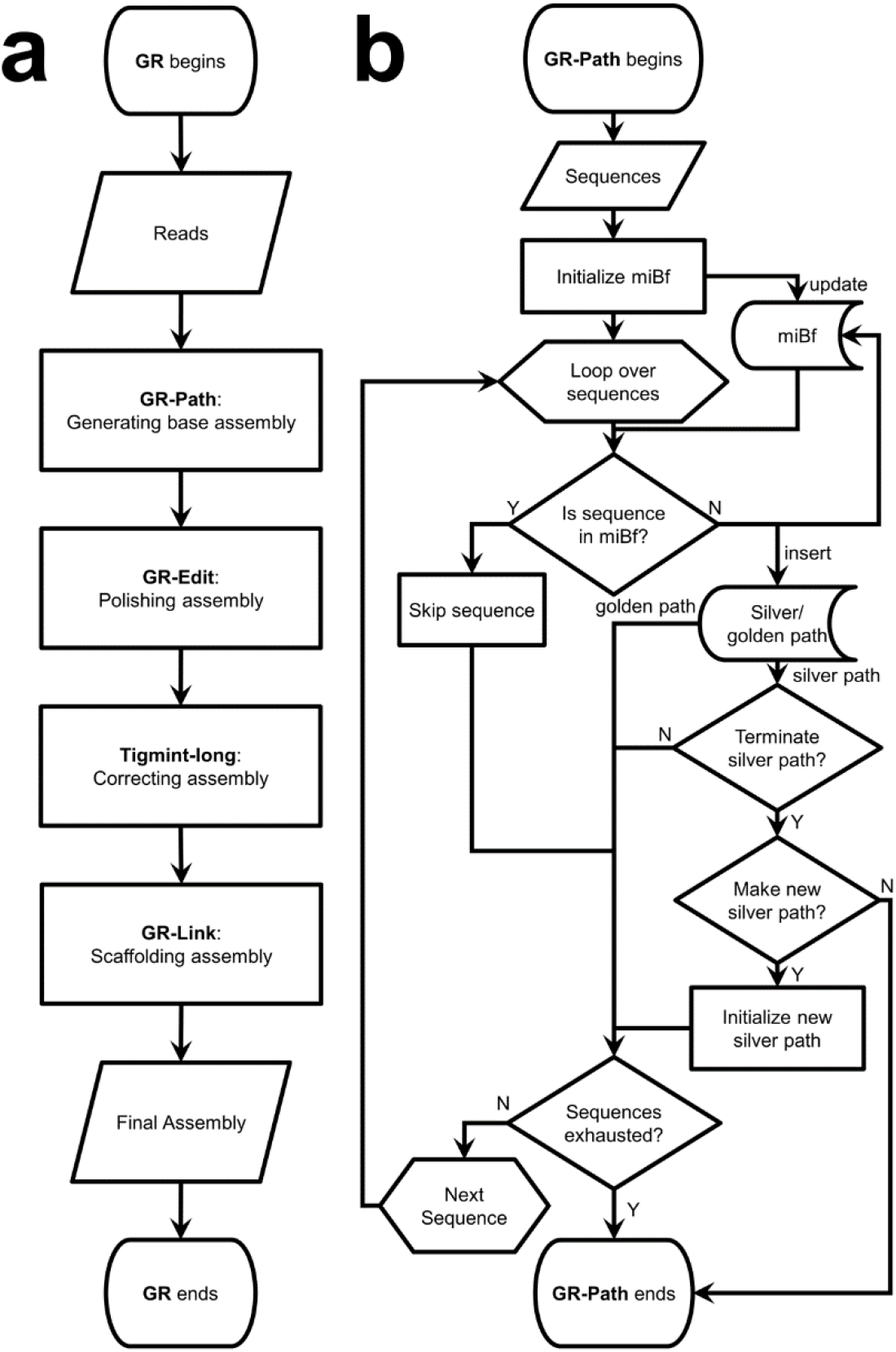
Flowchart of GoldRush (GR) and GoldRush-Path (GR-Path). In (a), raw long reads are first processed by GR-Path to generate the golden path, a ~1X representation of the genome. The golden path is then polished by GR-Edit and corrected for structural errors with Tigmint-long. Finally, GR-Link scaffolds the polished and corrected golden path to generate the final assembly. In (b), GR-Path uses the input long sequencing reads or silver path sequences to initialize a multi-index Bloom filter (miBf) data structure. GR-Path then loops over the sequences, and queries each read against the miBf. If the sequence is found in the miBf, GR-Path skips it and resumes its iterations. Conversely, if the sequence is not found in the miBf, it is inserted into the miBf and added to the silver/golden path. When GR-Path is constructing a silver path, if the silver path has not reached the threshold number of bases, GR-Path will continue recruiting bases from the input reads. If the threshold number of bases is reached, GR-Path will check if more silver paths need to be generated. If more silver paths are needed, GR-Path will create them using the same algorithm, otherwise it will terminate. Five (by default) silver paths, each representing ~0.9X (by default) coverage of the target genome, are combined to generate a low coverage subsample input for GR-Path to build the golden path. When creating the golden path, GR-Path will continue iterating over the sequences from the silver paths until all sequences are exhausted.

### 2.1 GoldRush-Path

The golden path is a ~1X representation of the target genome in read fragments and serves as the base assembly for the subsequent steps in GoldRush. Briefly, GR-Path iterates through the reads, querying each read against a data structure in turn, and inserts or skips over the read depending on the results of the query (Fig. 1b). Iteration over the long sequencing reads, as opposed to an all-vs-all alignment of reads, allows GoldRush to achieve a linear time complexity in the number of reads.

To create the golden path, GoldRush first builds silver paths. A silver path is similar to the golden path, except it is an ~*r*X (default 0.9) representation of the genome. The silver paths are combined to generate a low coverage subsample of the reads. This subsample is then used as input to GR-Path to generate the golden path.

GR-Path builds the silver paths by using a modified multi-index Bloom filter (miBf) (Chu et al., 2020), a resource-efficient probabilistic data structure, to associate spaced seed (a pattern with “care” and “don’t care” positions) derived k-mers with the locus of the genome they are derived from. The miBf is composed of three data structures: a Bloom filter, a rank array, and an ID array. The Bloom filter is first initialized with *h* (default 3) sets of spaced seed derived k-mers from the input read set using ntHash (Kazemi et al., 2022; Mohamadi et al., 2016). Only reads that are at least *m* bp (default 20,000 bp) long and have an average Phred quality score higher than *P* (default 15, representing 97% base accuracy or more, on average) are inserted into the Bloom filter. A rank array is then created to associate each set bit in the Bloom filter with a position in the empty ID array, sized based on the number of set bits in the Bloom filter.

The read set is then iteratively queried against the miBf data structure to determine if a read should be inserted into the miBf and thus the silver path. Once a read is processed and determined to contribute new base coverage to the silver path, the read is inserted into the miBf. The read is first split into tiles of length *t* (default 1,000), and *b* (default 10) consecutive tiles are binned into one block. The spaced seed derived k-mers from one block are hashed and a unique ID associated with the block, synonymous with a genomic locus, is inserted into the position of the miBf ID array that corresponds to the spaced seed derived k-mer (Supplementary Fig. S2). By binning tiles into blocks, GR-Path clusters together the spaced seed derived k-mers from a genomic region of length *b* × *t*. Each read that is inserted into the miBf is also saved to the silver path.

Like insertion, the querying process first splits the read into tiles of length *t*. The k-mers in these tiles are hashed using the same spaced seed patterns and queried against the miBf, then the associated ID hits within each tile are tallied in an ID-to-counts table. From these hits, a preliminary best ID hit is associated with each tile. The tiles are considered assigned (found) if the ID with the most hits exceeds a threshold *x* (default 10) or unassigned (not found) otherwise (Supplementary Fig. S3). Next, we improve the accuracy of the tile’s preliminary best hits by using information from the best hits of neighbouring tiles along with the target tile’s own ID-to-counts table (Supplementary Method S1, Supplementary Fig. S4).

Once GR-Path determines the final assignment of all the tiles in the read, it evaluates the read with three possible outcomes: skip, insert, or trim and insert. If all the tiles in the read are already assigned, the read is skipped since it does not contribute any new base information to the silver path. If all the tiles in the read are unassigned, the read is inserted into the miBf data structure and the silver path in its entirety. Finally, if the read has a mixture of assigned and unassigned tiles, the read is trimmed such that only one assigned tile on either side of the longest stretch of unassigned tiles is retained (Supplementary Fig. S5), and the trimmed read is inserted into both the miBf and silver path.

When the silver path contains or exceeds the threshold number of bases (default *genome size* × *r*), the current silver path is finalized, and a new silver path is initialized. The silver path creation continues until the input reads are exhausted or the number of completed silver paths reaches *M* (default 5). The golden path is then generated using the same algorithm and parameters used for the silver paths, except that the input is now the concatenated sequence files from the silver paths and the golden path is only complete when GR-Path has finished iterating through all of the silver path sequences. Sequences in the golden path construe the contigs of the initial assembly to be refined in the later steps of the pipeline, and are henceforth termed “goldtigs”.

### 2.2 GoldRush-Edit

After the golden path is built, the goldtigs are polished to correct mismatches and indels. The polishing protocol, GR-Edit, closely follows the ntEdit+Sealer protocol (J. X. Li et al., 2022; Paulino et al., 2015; Warren et al., 2019), which has been shown to perform well using short sequencing reads. Unlike the currently published paradigm, which stores all the short read k-mers in a single Bloom filter used for correction, GR-Edit uses a targeted approach where each goldtig has a dedicated Bloom filter, each containing a hash representation of k-mers derived from long read subsets.

To accomplish this, GR-Edit maps the long reads to goldtigs using, by default, minimap2 (H. Li, 2018). GR-Edit is also capable of using mappings from other tools, such as ntLink (Coombe et al., 2021). For each goldtig, the set of mapped reads are k-merized using a range of different k-mer lengths in order to benefit from a trade-off between specificity (longer k-mers) and sensitivity (shorter k-mers). These k-mers are then inserted into an array of Bloom filters, one Bloom filter for each k-mer size per goldtig. From this, we have a set of k-mers targeted to individual goldtigs that we can use with ntEdit and Sealer for polishing (Supplementary Fig. S6).

GR-Edit then polishes the goldtigs using the full ntEdit+Sealer pipeline on each individual goldtig using their dedicated Bloom filters. GR-Edit launches multiple ntEdit+Sealer pipelines in parallel to amortize the overhead introduced with each polishing run.

### 2.3 GoldRush-Link

After polishing and correcting the goldtigs, an updated version of the long-read scaffolder ntLink (Coombe et al., 2021) is used to assemble the goldtigs. To utilize the long read evidence in building longer sequences, the full long read set is mapped to the goldtigs using a lightweight minimizer-based approach. Briefly, minimizer sketches are generated for the goldtigs as well as each read for a given k-mer size *k* and window size *w*, as described previously (Coombe et al., 2021). The goldtig minimizer sketches are indexed, and for each minimizer in the sketch of a given long read, this index is queried to find hits between the long read and the goldtigs. Long read mappings that span multiple goldtigs provide scaffolding evidence. This long read evidence is stored as a scaffold graph, where the nodes are goldtigs, and the directed edges between the nodes represent evidence that the goldtigs should be joined. This scaffold graph is traversed using abyss-scaffold to output the final, contiguated assembly (Jackman et al., 2017).

Three important new features have been added to ntLink (v1.3.0+) to adapt the functionality for the *de novo* long read assembly problem in GoldRush: overlap detection, gap-filling, and scaffolding rounds based on liftover of sequence mappings.

For the sequence pairs with putative overlaps, minimizer sketches are generated with a lower *k* and *w* than the initial ntLink pairing stage to increase the sensitivity of overlap detection (parameters *small_k, small_w*, defaults 15 and 10, respectively). These sketches are filtered to retain minimizers that fall in the estimated overlapping region with a multiplicity of one in each sequence, ensuring that only non-repetitive minimizers in the sequences are retained. These minimizers are then used to create an undirected minimizer graph, similar to methods employed by the reference-guided scaffolder ntJoin (Coombe et al., 2020). In this graph, the minimizers are nodes, and edges between the minimizers indicate that the minimizers are adjacent in at least one of the ordered sequence minimizer sketches, with the edge weights indicating the number of sequences that have that minimizer adjacency. This minimizer graph is filtered to retain edges with a weight of 2, which removes branches and results in a graph consisting of linear path components. Each linear path is a minimizer-based mapping between the putatively overlapping sequence ends. The middle minimizer from the longest mapping is chosen to anchor the sequences to one another, and the coordinates of this minimizer guide the trimming of the detected overlapping regions on the incident sequences. Finally, after this trimming, the sequences are concatenated (Supplementary Fig. S7).

The second major feature added to ntLink uses the mapped long reads to fill gaps between the scaffolded goldtigs. The verbose option in the initial ntLink pairing stage was updated to output the complete long read mapping information, including the mapped goldtigs and the minimizers (including position and strand on the goldtig and read). For each sequence join induced by ntLink, the verbose mapping information is parsed to identify each read that supports that join, and the associated mapped minimizers (“pass 1 minimizers”). The read with the highest average number of mapped pass 1 minimizers is chosen and subsequently used to fill the scaffold gap. As finding anchoring minimizers as close to the sequence ends as possible is preferred, each chosen read is re-mapped to the flanking sequences using a lower k and w for increased sensitivity (“pass 2 minimizers”). If the mapping is unambiguous, the anchoring pass 2 minimizers closest to the sequence ends are used as cut points for the flanking sequences and the read sequence filling the gap. Otherwise, the pass 1 minimizers are used to determine the gap-filling coordinates (Supplementary Fig. S8). Gap-filling is turned on in GoldRush by default and is run when the target “gap_fill” is specified to the ntLink command.

Finally, liftover-based rounds were integrated into the ntLink code base. We added a step to liftover the mapped minimizer coordinates in the verbose mapping file (described above) from the initial goldtigs to the sequences post-scaffolding. This new mapping file is then input to the ntLink pairing stage in the subsequent ntLink round, which uses the input mapping coordinates instead of re-mapping the reads to infer the scaffold graph. The remaining steps in the ntLink pipeline then proceed as previously described. To invoke these liftover-based rounds, we provided a new Makefile “ntLink_rounds”, which runs a specified number of rounds of ntLink (parameter *rounds*, default 5), lifting over the mapping coordinates between each iteration.

### 2.4 Implementation

The GoldRush pipeline is driven by a Makefile. GR-Path and GR-Edit are coded in C++, and GR-Link is coded in Python, using the btllib library (Nikolić et al., 2022). The tool can be installed from GitHub or using the conda package manager. Instructions on how to run the GoldRush pipeline are provided on the GitHub page (https://github.com/bcgsc/goldrush). Many of the GoldRush parameters are supplied with default values and can be configured. Only the genome size of the target species and the long reads in a single, uncompressed, multi-FASTQ file are required as input.

### 2.5 Evaluation

To evaluate the performance of GoldRush (v1.0.0), we assembled five genomes from ONT long-read data for three human cell lines (NA24385, HG01243, and HG02055), *Oryza sativa* (rice), and *Solanum lycopersicum* (tomato) (Supplementary Table S1). We optimized the parameters of GoldRush for each dataset (Supplementary Table S2, Supplementary Figs S9-S13). In separate trials, we also polished the golden paths with Racon (v1.5.0) (Vaser et al., 2017) instead of GR-Edit. To compare the performance of GoldRush to current state-of-the-art long read assemblers, we assembled all five datasets with Flye, Redbean, and Shasta. We ran both Flye (v2.9) and Redbean (v2.5) using their default parameters, and Shasta (v0.10.0) using the “Nanopore-Plants-Apr2021.conf” configuration file for *O. sativa* and “Nanopore-May2022.conf” for the other datasets.

All assemblies were analyzed using QUAST (v5.0.2) (Gurevich et al., 2013) (--fast --large --scaffold-gap-max-size 100000 --min-identity 80 --split-scaffold), and the corresponding reference genome (Supplementary Table S3). To assess the contiguity and correctness of the assemblies, we report the NG50 and NGA50 length metrics, and the number of extensive misassemblies (as defined by QUAST). The NG50 length statistic describes that 50% of the genome size is in sequences the NG50 length and longer. The NGA50 length statistic is similar to the NG50 length, but uses alignment blocks instead of sequence lengths for the calculation. We also ran BUSCO (v5.3.2) (Simão et al., 2015) using the primates_odb10 lineage to assess the completeness of the human assemblies in the gene space. All benchmarking tests were performed on a server-class system with 144 Intel(R) Xeon(R) Gold 6254 CPU @ 3.1 GHz with 2.9 TB RAM.

## 3 Results

We assembled the genomes of three different human cell lines (NA24385, HG01243 and HG02055), *O. sativa*, and *S. lycopersicum* using GoldRush, Flye, Redbean, and Shasta, and compared the resulting assemblies using a variety of length contiguity metrics, assembly accuracy markers, including those reported by QUAST (Gurevich et al., 2013) and BUSCO (Simão et al., 2015), and their resource usage.

### 3.1 Assembly Performance

#### 3.1.1 Contiguity and Correctness

For the genome assemblies of all three human cell lines, GoldRush achieved NG50 lengths between 25.3 and 32.6 Mbp, comparable to both Shasta (29.7-39.6 Mbp) and Flye (26.6-38.8 Mbp), and typically 3 times more contiguous than the Redbean assemblies (8.0-10.9 Mbp) (Supplementary Tables S4-S6). Two of the three human GoldRush genome assemblies (NA24385 and HG01243) also had the fewest extensive misassemblies (940 and 1,057) among the tools tested; ~2-3 times fewer than Shasta (1,682 and 3,240), and ~5-7 times fewer than Redbean (4,918 and 7,052). Despite the relatively low number of structural misassemblies, the NGA50 length for each human GoldRush assembly is around 20 Mbp, indicating that some misassemblies are found in the larger scaffolds and are breaking the alignment blocks (Fig. 2a). In addition to assembling highly contiguous human genomes, GoldRush is also robust in assembling plant genomes, reaching 0.3 Mbp and 2.5 Mbp NGA50 lengths for *O. sativa* and *S. lycopersicum*, respectively (Fig. 2c, Supplementary Tables S7 and S8). The *O. sativa* Shasta genome assembly, on the other hand, had scaffold NG50 and NGA50 lengths of 124,700 bp and 104,593 bp, respectively, only ~4.2 and ~3.6 times longer than the raw reads used as input for assembly, respectively (N50=29,349 bp) (Supplementary Tables S1 and S8).

**Fig. 2.**
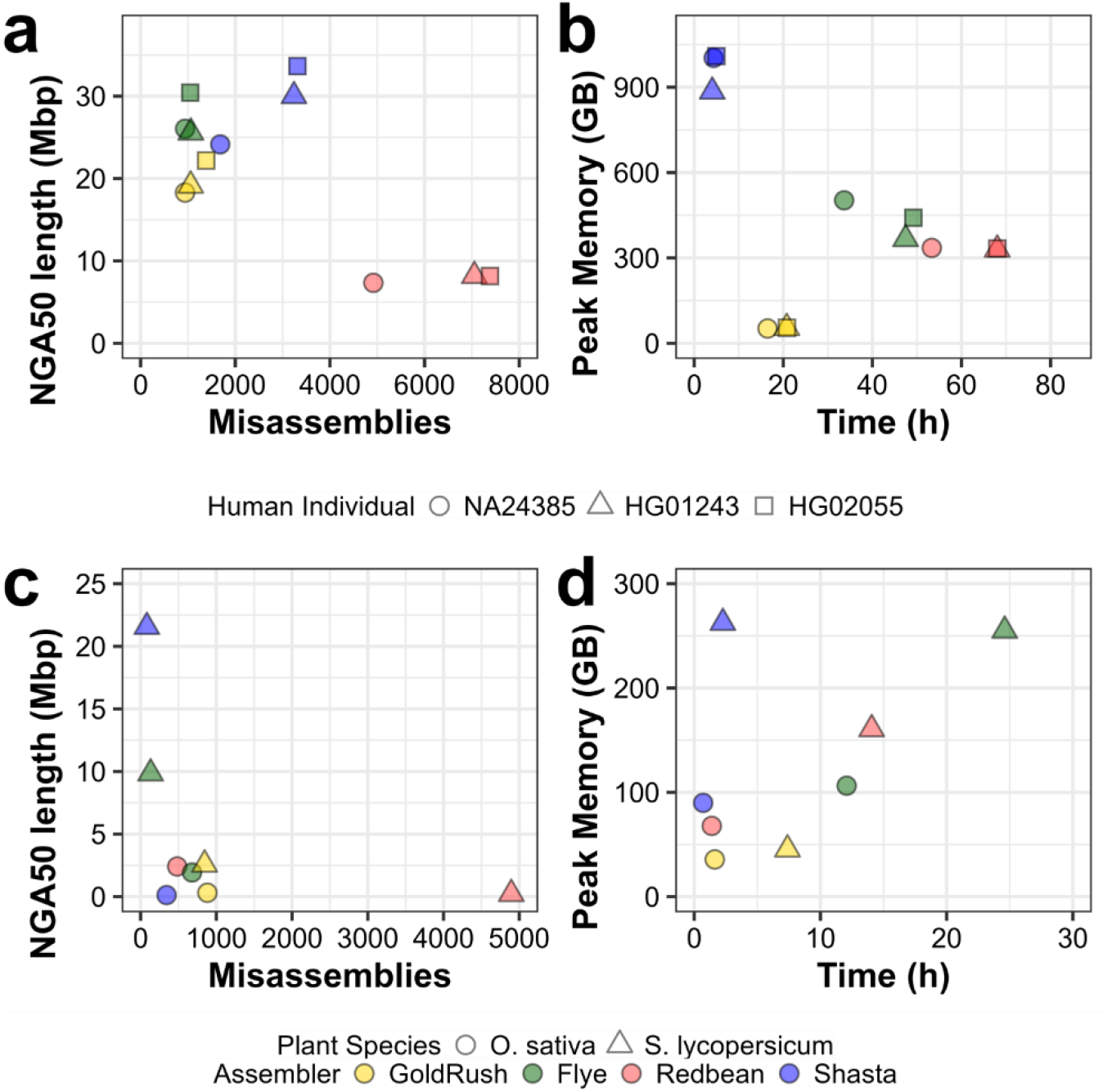
Contiguity, correctness, and resource usage of GoldRush assemblies compared to Flye, Redbean, and Shasta assemblies for three human individuals (NA24385, HG01243, and HG02055), *O. sativa*, and *S. lycopersicum*. The genome assemblies were assessed using QUAST for their contiguity and correctness. Extensive misassemblies and NGA50 length as determined by QUAST are shown on the x and y axes, respectively, in (a) and (c). Peak memory (GB) usage and runtime (h) of the assembly processes were recorded using the unix time command and shown on the x and y axes, respectively, in (c) and (d).

#### 3.1.2 Computational Performance

GoldRush and Shasta assembled all three human genomes in less than 24 h (Fig. 2b). Both Flye and Redbean required at least a day to assemble each of the three human genomes, with Redbean assembling two of the datasets (HG01243 and HG02055) in 68 h, for each (Fig. 2b, Supplementary Tables S9-S11). GoldRush is also competitive in assembling the smaller plant genomes, requiring less than 2 and 8 h to assemble *O. sativa* and *S. lycopersicum*, respectively (Fig. 2d, Supplementary Tables S12 and S13). GoldRush used at most 54.5 GB of RAM to assemble the three human genomes (Fig. 2b, Supplementary Tables S9-S11). In comparison, using the same data, Flye and Redbean used between 329.3-502.4 GB (6 to 8-fold more than GoldRush), and Shasta utilized 884.8-1,009.2 GB (up to 20-fold more than GoldRush). Similarly, GoldRush required the least amount of RAM to assemble the *O. sativa* and *S. lycopersicum* datasets, using at most 45.3 GB (Fig. 2d, Supplementary Tables S12 and S13).

### 3.2 GoldRush-Edit Polishing

GR-Edit decreased the number of mismatches and indels per 100 kbp of the NA24385 golden path by ~6.5 fold (1,463.7 to 228.5 and 1,327.2 to 197.2, respectively) (Supplementary Table S14). This improvement in mismatches and indels translated into a recovery of 12,272 (89.1%) complete BUSCOs, fewer than the 12,920 (93.8%) and 12,988 (94.3%) complete BUSCOs in the Shasta and Flye assemblies, respectively, but more than the 12,193 (88.5%) complete BUSCOs reconstructed in the Redbean assembly (Supplementary Table S15). The results of running BUSCO on the HG01243 and HG02055 genome assemblies can be found in Supplementary Tables S16 and S17. When substituting GR-Edit with Racon for polishing the same golden path, the polishing step of the GoldRush pipeline incurred a greater computational cost, requiring over an order of magnitude more memory (602.3 vs 11.0 GB RAM) but resulted in a more base-accurate assembly (157.0 and 106.4 mismatches per and indels per 100 kbp, respectively) (Fig. 3, Supplementary Table S18). The improvements in the base accuracy of the resulting NA24385 genome assembly also translated in a higher recovery of complete BUSCOs (12,752, 92.6% complete) (Supplementary Table S19).

**Fig. 3.**
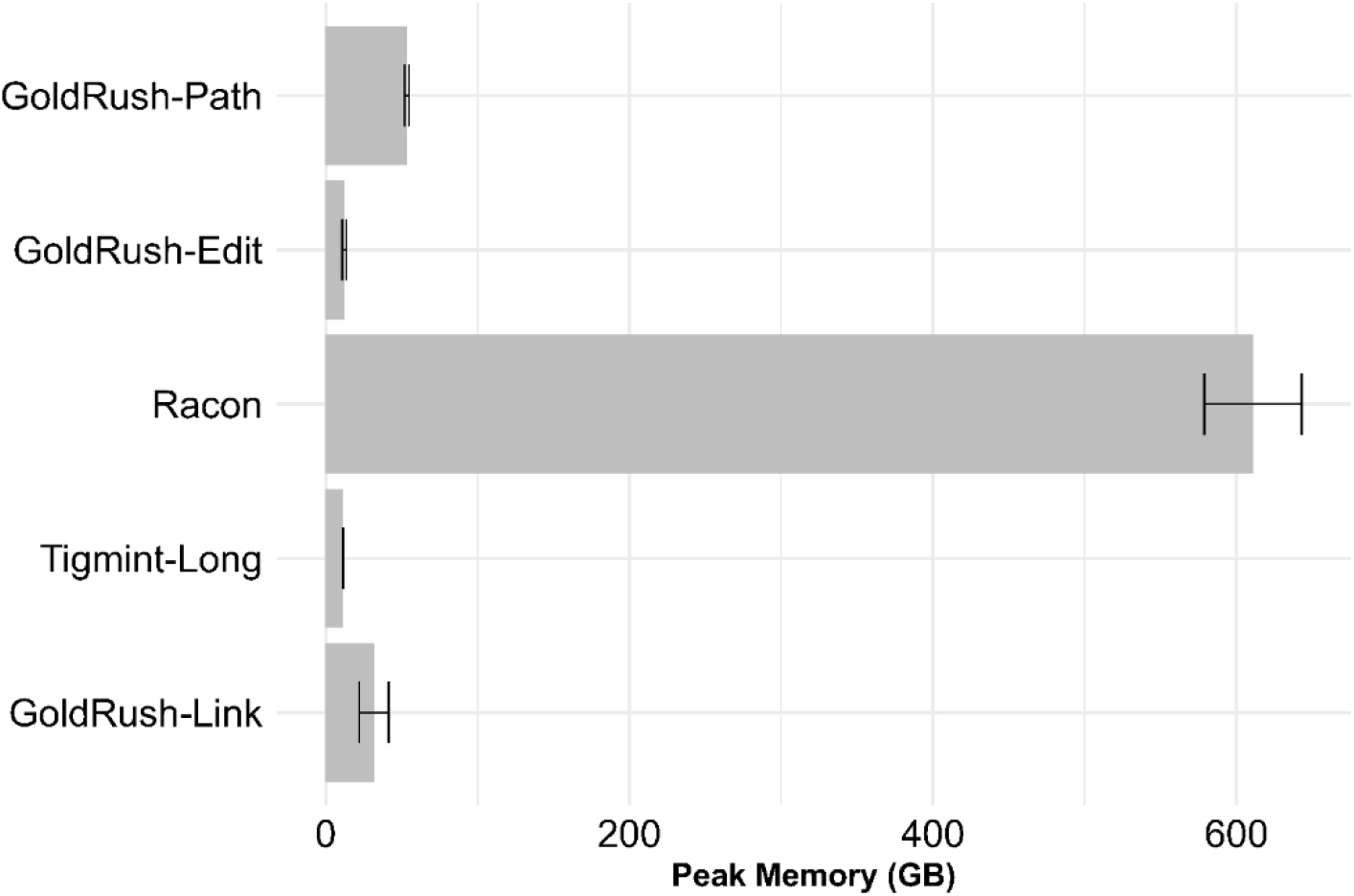
Memory usage of the GoldRush stages when assembling human genome datasets (individuals NA24385, HG01243, and HG02055). The average peak memory for the three human genomes is shown, with the error bars indicating the standard deviation. Racon can optionally be used for long read base polishing within GoldRush, if the onboard memory of the computer system is not limited.

## 4 Discussion

The GoldRush algorithm is straightforward: collect unique fragments representing the genome to generate a golden path, polish the fragments, correct them for structural misassemblies, and join the polished and corrected fragments together. As GoldRush is built upon this fundamental concept of the golden path, it represents a paradigm shift in the assembly of erroneous long reads, no longer requiring the time- and memory-intensive process of all-vs-all alignments. Instead, the golden path, or a ~1X read fragment representation of the underlying genome, is constructed through iterating over the read set, and querying a progressive miBf database representing the golden path. We have shown that this novel assembly paradigm yields human genome assemblies that are comparable in contiguity to what can be obtained using different implementations of the OLC algorithm, yet with an order of magnitude smaller memory footprint.

The GoldRush algorithm was designed with no single long-read sequencing technology in mind, making it versatile and platform agnostic. The algorithm is robust to base errors, capable of assembling long read datasets with estimated error rates ranging from 4% to 20%, and achieving NG50 and NGA50 lengths up to 32.6 and 22.2 Mbp, respectively, for the human data tested (Supplementary Tables S1, S4-S8). GoldRush accomplishes this by mitigating the impact of base errors at various stages. GR-Path uses spaced seeds to enable more sensitive detection of erroneous sequences originating from the same genomic locus. In addition, GR-Path only selects sequences with Phred qualities >= 15 (by default) for the silver and golden paths. This requirement ensures that the baseline assembly is composed of the higher quality reads from the set. For polishing the golden path sequences GR-Edit accommodates the higher error rate of long reads by utilizing targeted Bloom filters populated with localized k-mers originating exclusively from mapped reads. This approach of using a targeted Bloom filter per goldtig enables the use of smaller k-mer sizes than those used for the original short-read focused ntEdit+Sealer protocol, thus increasing the polishing sensitivity and mitigating errors that would otherwise arise from the use of off-target k-mers. Since GoldRush uses a completely different paradigm to assemble genomes, it also has different strengths when compared to OLC-based genome assemblers. For instance, in the *O. sativa* dataset the N50 length of the reads is 29,349 bp, and a Shasta assembly of the data results in an assembly NGA50 length of 104,593 bp – roughly 3.6 times longer. In contrast, the NGA50 length of the GoldRush assembly is 10-fold greater (307.9 kbp) than the read N50 length (Fig. 2c, Supplementary Tables S1 and S8). However, when considering which tool performs best on all datasets, none consistently outperforms the other in all the metrics we measured (i.e. run time, memory usage, genome contiguity, genome completeness, and genome correctness).

Yet, GoldRush is consistently more memory efficient in comparison to other tools in all the datasets tested. This provides the opportunity to assemble long read data from human-sized or larger genomes to those who do not have access to server-class systems, especially as retail computers with 64 GB RAM or more become more accessible. The memory efficiency of GoldRush is mainly due to the use of the miBf data structure in the GR-Path stage. However, the published version of the miBf data structure (Chu et al., 2020) was intended to serve as a static database, where the user first inserted all the items of interest, and subsequently used the database solely for query operations. For the purpose of GR-Path, we needed a memory-efficient data structure that could also be dynamic, with interleaved insert and query operations. To adjust the miBf for GR-Path, we disabled its ability to rescue information lost to hash collisions, a step that requires all the elements to be inserted and remain static afterwards. We compensated for this loss of information by using longer tile lengths, *t*. With longer tiles, there are more queries per given tile, and the increased number of queries would offset the loss of expected hits due to hash collisions.

GoldRush also assembles all the human datasets within a day and, together with Shasta, is faster than all other OLC assemblers tested herein. Shasta accomplishes this with heuristics based on MinHash markers to quickly identify potential read-to-read overlaps and speed up the overlap component of OLC (Shafin et al., 2020). Breaking down the time GoldRush spends for completing each stage, we observe that GoldRush devotes more time polishing the golden path with GR-Edit, which is already heavily optimized (Supplementary Table S20-S22). GR-Edit runs a background Bloom filter building process which continually produces Bloom filters from the mapped reads for the launched pipelines, minimizing waiting time between ending and starting a new polishing run (Supplementary Fig. S14). The GR-Path and GR-Link stages were also optimized to reduce the overall runtime of GoldRush. GR-Path utilizes multiple independent silver paths to effectively generate a low coverage subsample of the original dataset, such that the subset still covers the entire target genome. With enough silver paths, any sections of the genome that are missing in a given silver path should be recovered in the others. This enables GR-Path to generate a golden path without having to process the entire dataset, speeding up the golden path generation considerably.

Finally, we observed that running additional rounds of ntLink in the GR-Link stage led to substantial improvements in the contiguity of the final assembly. However, re-mapping the long reads for each round was costly. To remedy this, we implemented a mapping liftover step, allowing ntLink to run multiple rounds of scaffolding without re-mapping the long reads in each round. We also implemented two additional features, overlap detection and gap-filling, in ntLink. While these were introduced specifically for the GR-Link stage of GoldRush, they are also applicable to the general use of ntLink. In earlier versions of ntLink, overlapping sequences were still concatenated end-to-end, which could result in local insertion misassemblies at the contig joints. With the new overlap detection and overlap resolution feature, gap estimates from the earlier stages of ntLink are used to identify putative overlaps between adjacent goldtigs (indicated by negative gap estimates), and guide trimming of the overlapping sections. Overlapping goldtigs are expected in the golden path, as reads are evaluated on a tile-by-tile basis in GR-Path. Furthermore, the new gap-filling feature can help to recover missing sequence in the golden path reconstruction if there are gaps between adjacent golden path sequences.

With the recent release of the ONT Q20+ chemistry and its purported base accuracy of 99%, as well as the continual improvements in basecallers, we expect GoldRush to capitalize on the improvements in sequencing accuracy and reduce the time spent correcting base errors. Indeed, for the human GoldRush tests, GR-Edit executed the fastest for the NA24385 dataset, which is estimated to be the least erroneous (4%) (Supplementary Tables S1, S20-S22). Lastly, GoldRush is modular. Each step within GoldRush can be substituted for another tool that performs the equivalent function, such as substituting GR-Edit for Racon, allowing GoldRush to easily benefit from any future advances in the field. GoldRush also makes no assumptions about the quality of the input long reads, standing only to gain from future computing and sequencing improvements in the long read sequencing domain.

We have demonstrated that our memory-efficient and modular long read assembly pipeline, GoldRush, assembles long ONT reads into draft genomes with high contiguity, notably chromosome 18 telomere-to-telomere (HG02055) and several other chromosome arms (Supplementary Fig. S15). We also show the assembly contiguities are comparable to what is achieved with current state-of-the art tools, but GoldRush uses a fraction of the RAM, lowering the barrier to entry to human long read assembly. With its modular design, memory-efficiency, and robust performance in assembling large and complex genomes, we envision GoldRush to be beneficial to the scientific community and expand the reach of long sequencing reads.

