## Supplementary Material for "GoldRush: A *de novo* long read genome assembler with linear time complexity"

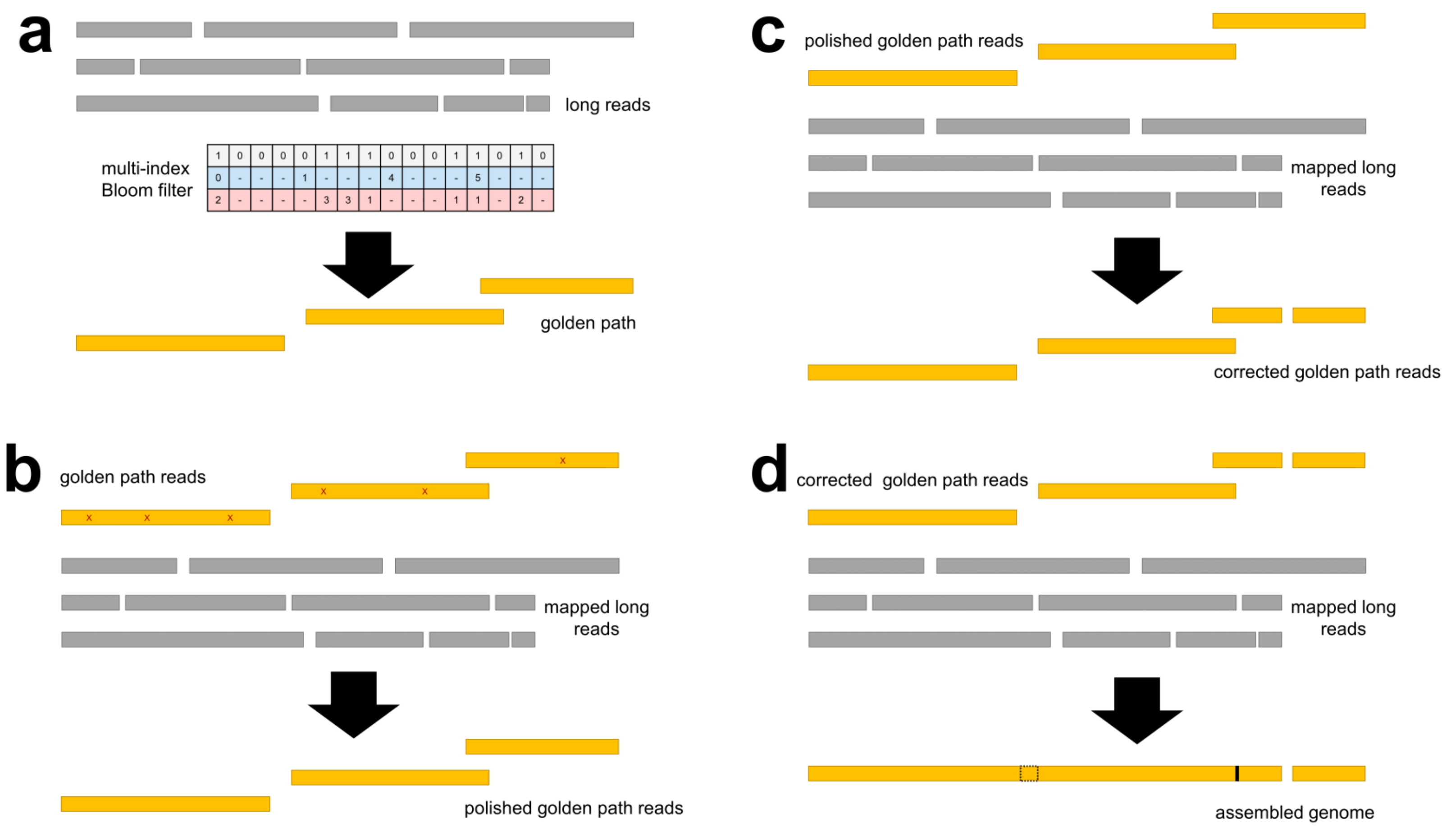

**Supplementary Fig. S1. Flowchart of the GoldRush assembly pipeline.** In (a), GR-Path queries the long reads against the multi-index Bloom filter to generate the golden path, a ~1X representation of the underlying genome. The golden path (errors shown as red "X"s) is then polished by GR-Edit in (b), and corrected with Tigmint-long in (c). Finally, GR-Link scaffolds (gap filling shown in dotted box and trimming shown in black) the polished and corrected golden path to generate the final assembly in (d).

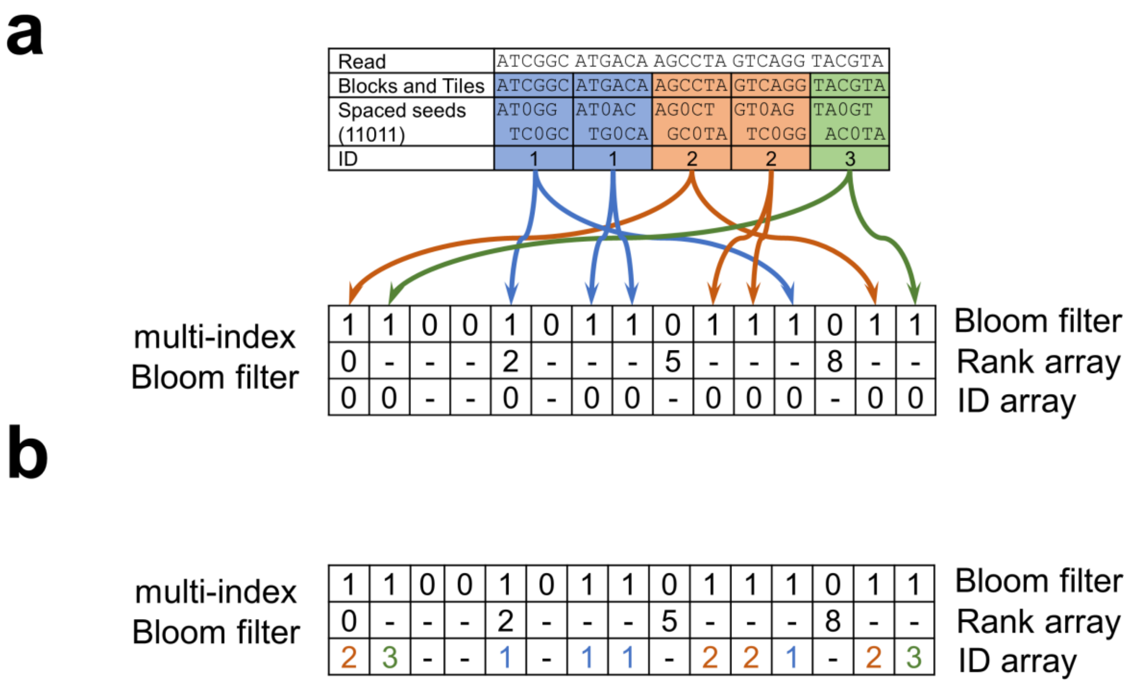

**Supplementary Fig. S2. Example of logic for inserting read signatures to the miBf data structure and golden path.** A read is being inserted into the miBf in (a). The read is inserted using *t* (length of tile) = 6, *b* (number of tiles in a block) = 2, *h* (number of spaced seed patterns) = 1, with a spaced seed of pattern of 11011. The read is first divided into tiles of length *t*. This is represented by each smaller cell. Cells that belong to the same bin are given the same colour. The blue, orange, and green cells are associated with the IDs 1, 2, 3, respectively. The sequence in each tile is hashed using the spaced seed pattern and inserted into the miBf. (b) shows the changes in the miBf after insertion. The colors of the IDs in the ID array in (a) correspond to the blocks they are derived from in (a).

**a**

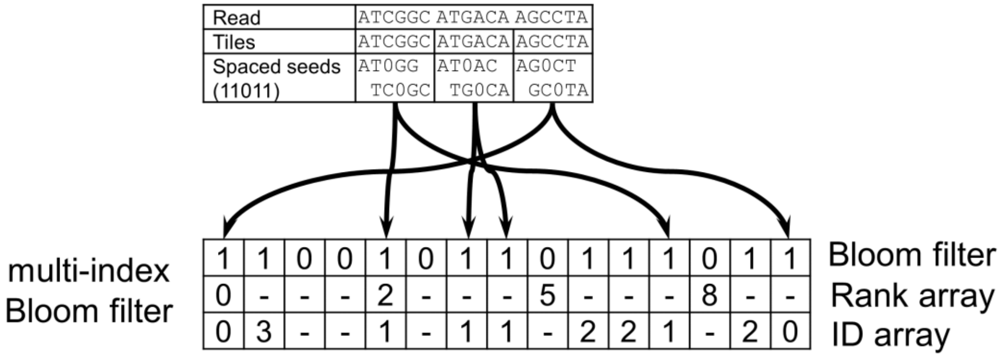

**b**

| | | | |
|---|---|---|---|
| Read | ATCGGC ATGACA AGCCTA | | |
| Tiles | ATCGGC | ATGACA | AGCCTA |
| Spaced seeds (11011) | AT0GG TC0GC | AT0AC TG0CA | AG0CT GC0TA |
| ID: counts tally | 1:2 | 1:2 | |
| Associated ID | 1 | 1 | 0 |
| Tile classification | true | true | false |

**Supplementary Fig. S3. Example of querying the miBf with a new read.** A read is being queried against the miBf in (a). The read is queried using *t* (length of tile) = 6, and *x* (hit threshold for a tile to be assigned) = 1, with a spaced seed of pattern of 11011. The read is first divided into tiles of length *t*. This is represented by each smaller cell. The sequence in each tile is hashed using the spaced seed pattern and queried against the miBf. (b) shows the results of each cell's query. Both the first and the second cell have two hits to ID 1. GR-Path will then associate these two tiles with and ID of 1 assess these two tiles as true (assigned) because the number of hits to ID 1 is greater than x, which is 1. On the other hand, the are no hits for the third tile. GR-Path will associate the tile with an ID of 0, assess it as false (unassigned).

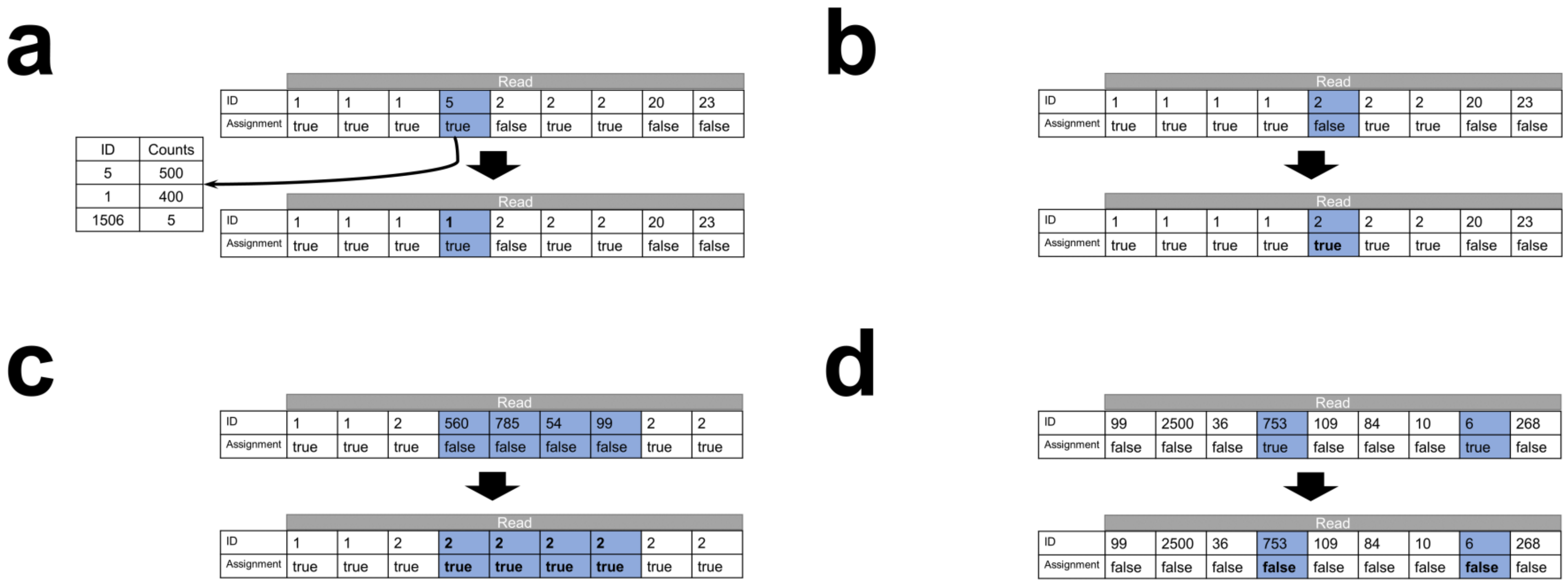

**Supplementary Fig. S4. Improving the accuracy of the tile's best hit.** In (a), GR-Path identifies the tile with an associated of ID of 5 for improvement (shaded in blue). The neighbour IDs are 1 and 2, and there is an entry in the ID-to-count tables with an ID of 1, so the tile's ID is set to 1. In (b), GR-Path identifies the unassigned tile with an associated of ID of 2 for improvement (shaded in blue). The adjacent tiles have the same ID or 1 smaller than the current tile, so the current tile is set as assigned. In (c), a stretch of unassigned tiles is flanked by assigned tiles with the same ID (shaded in blue). The stretch of unassigned tiles is then set to assigned and have their IDs changed to 2. In (d), there are isolated assigned tiles flanked by unassigned tiles (shaded in blue). These are likely tiles assigned due to false positive hits, so these tiles are set to unassigned.

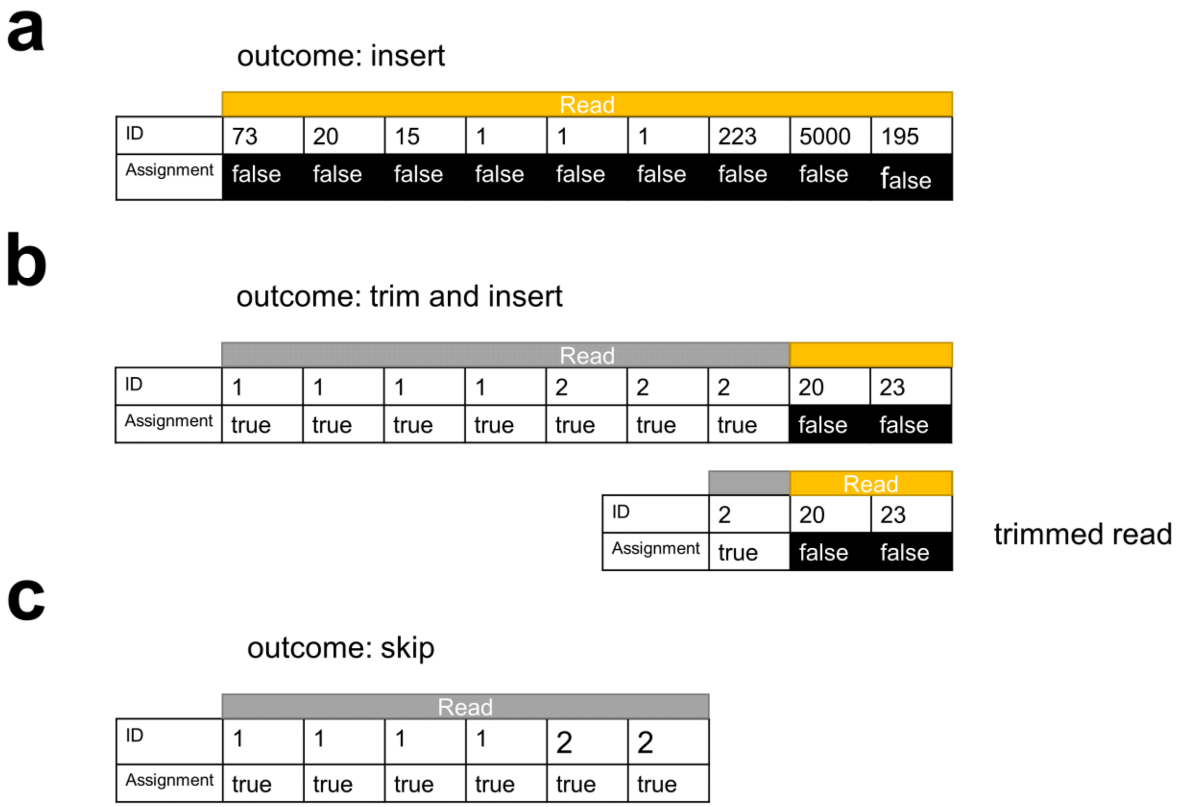

**Supplementary Fig. S5. Example of the three different outcomes when querying a read.** In (a), every tile of the read is unassigned (false) which means the genomic locus the read is sequenced from is not captured in the miBf, so we insert the read into the miBf. In (b), the read has a mixture of assigned and unassigned tiles. In this case, we trim the read and retain one tile overhang from the unassigned region. The trimmed read is inserted into the miBf. In (c), every tile in the read is classified as assigned. Therefore, the genomic locus the read represents is already captured in the miBf and golden path, and the read is skipped.

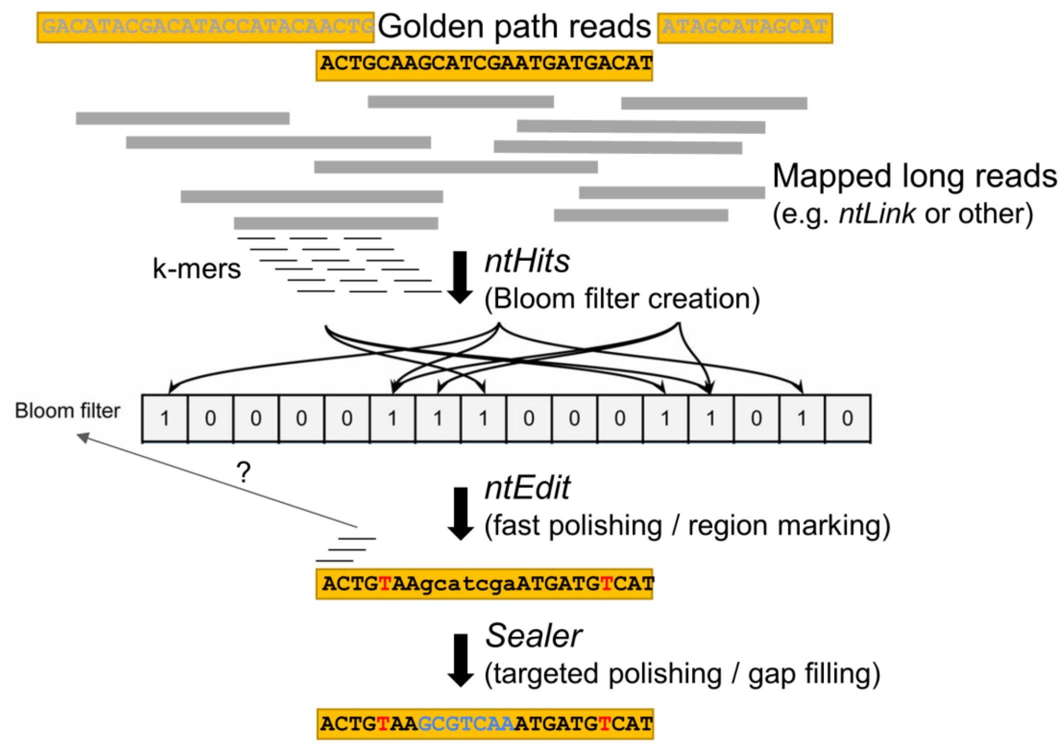

**Supplementary Fig. S6. Schematic describing the GoldRush-Edit polishing protocol.** Mapped long reads are k-merized and inserted into an array of targeted Bloom filters with different k-mer sizes using ntHits. Then, ntEdit uses these Bloom filters to correct mismatches and small indels, and marks the regions it is unable to fix. Finally, Sealer detects these unresolved regions as gaps and attempts to fill them using the targeted Bloom filters.

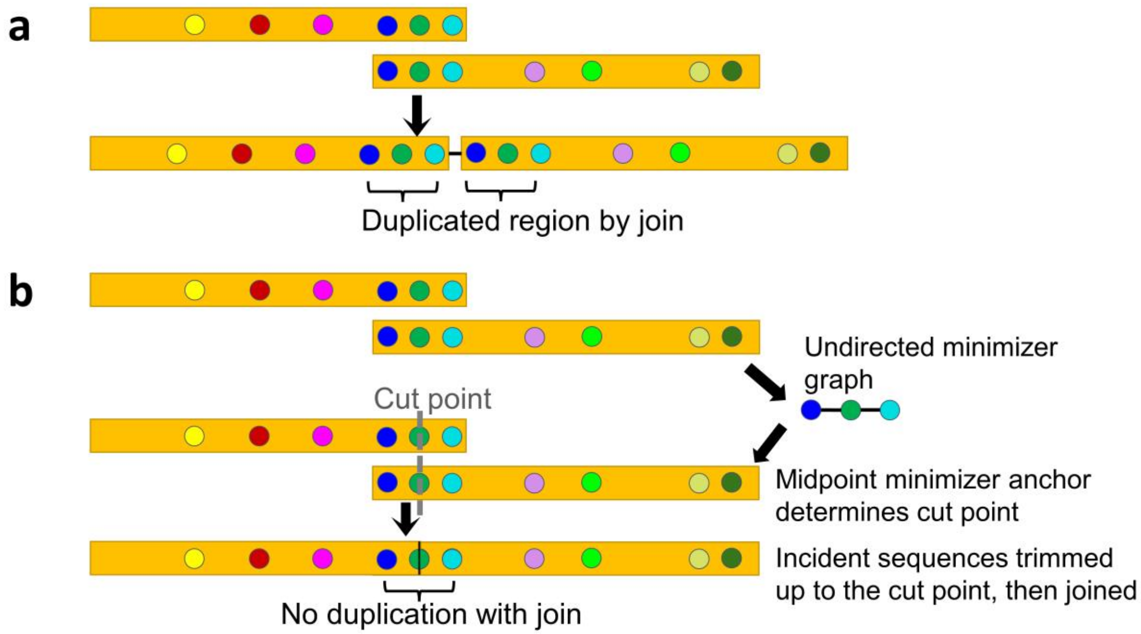

**Supplementary Fig. S7. The overlap detection and resolution feature of ntLink.** In these schematics, the circles represent minimizers, and the same colour indicates the same minimizer. (a) In previous versions of ntLink, overlapping sequences were joined end-to-end, which created small insertion misassemblies. (b) The overlap detection and resolution mode developed in ntLink detects and resolves these overlapping regions to avoid these spurious insertions. First, minimizers with a smaller *k* and *w* compared to the ntLink pairing stage are computed in the putative overlap region for a pair of sequences. The shared minimizers in this region with a multiplicity of one in each sequence (dark blue, medium green and light blue minimizers in (b)) are retained, and used to create an undirected minimizer graph. In this graph, the nodes are minimizers and adjacent minimizers are connected by edges. This graph is traversed to detect a mapping block between the sequence ends, and the position of the middle minimizer in the block chosen as the cut point. Finally, each sequence is trimmed up to the cut point and concatenated.

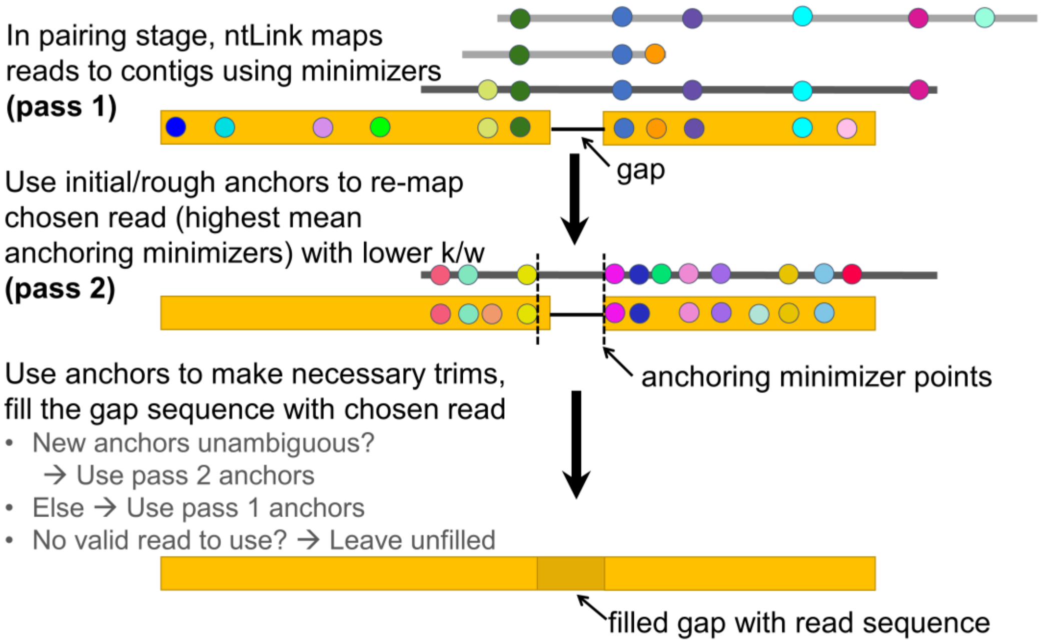

**Supplementary Fig. S8. The gap-filling feature of ntLink.** In filling a gap between a sequence pair, the read that supports the join with the highest mean number of anchoring minimizers is chosen as the representative sequence to fill the gap (pass 1 minimizers). The read is re-mapped to the incident sequences with a lower k and w for a more specific mapping (pass 2 minimizers). If the pass 2 minimizers are unambiguous, the anchoring minimizers are determined with these mappings, otherwise the pass 1 minimizers are used. These anchoring minimizers guide the trimming of the incident sequences as well as the chosen read for gap-filling. Finally, the trimmed sequences are concatenated together to output the scaffold with the filled gap.

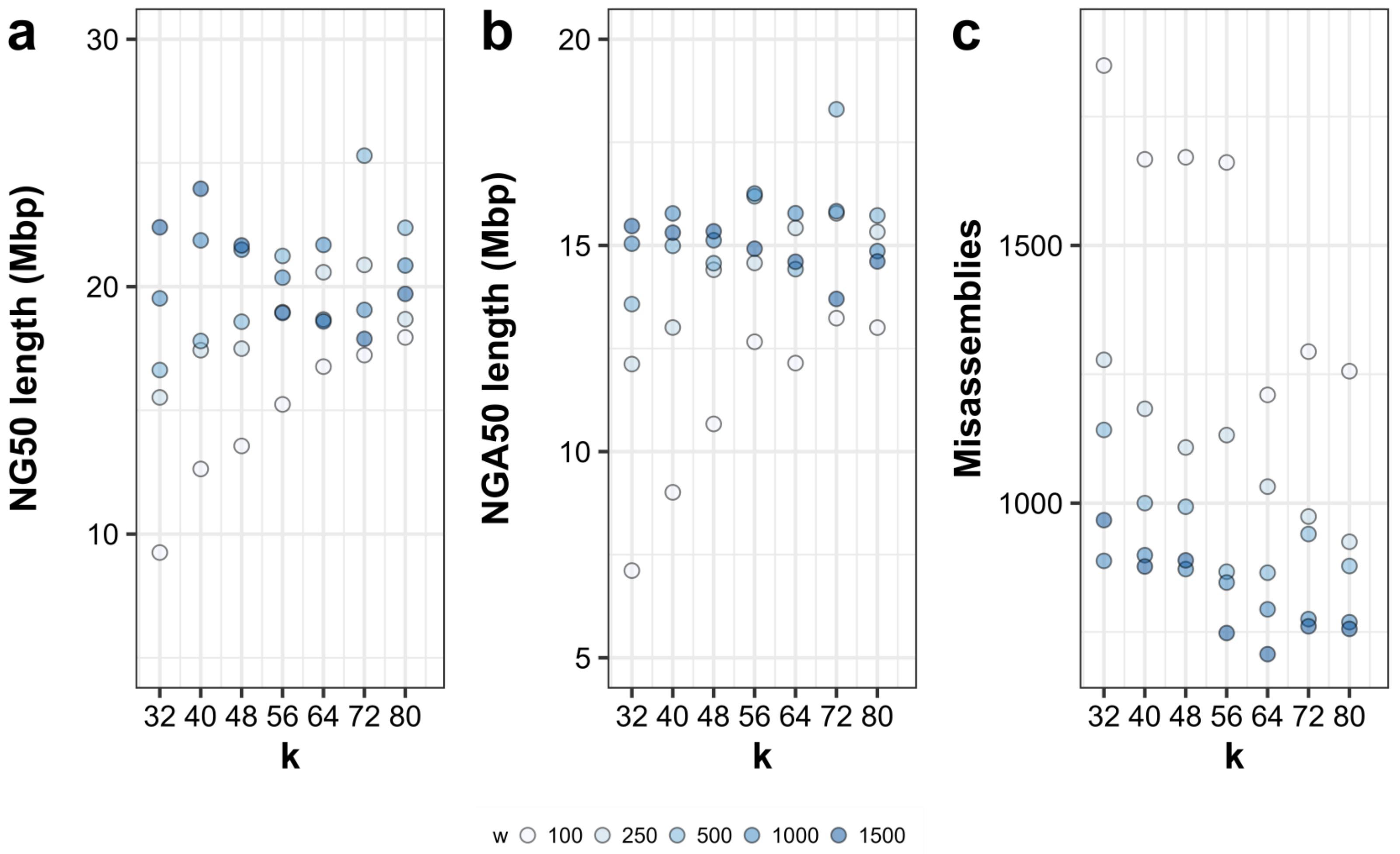

**Supplementary Fig. S9. Contiguity and correctness results of assembling long reads from the human individual NA24385 with GoldRush, sweeping on the GoldRush-Link k and w parameters.**

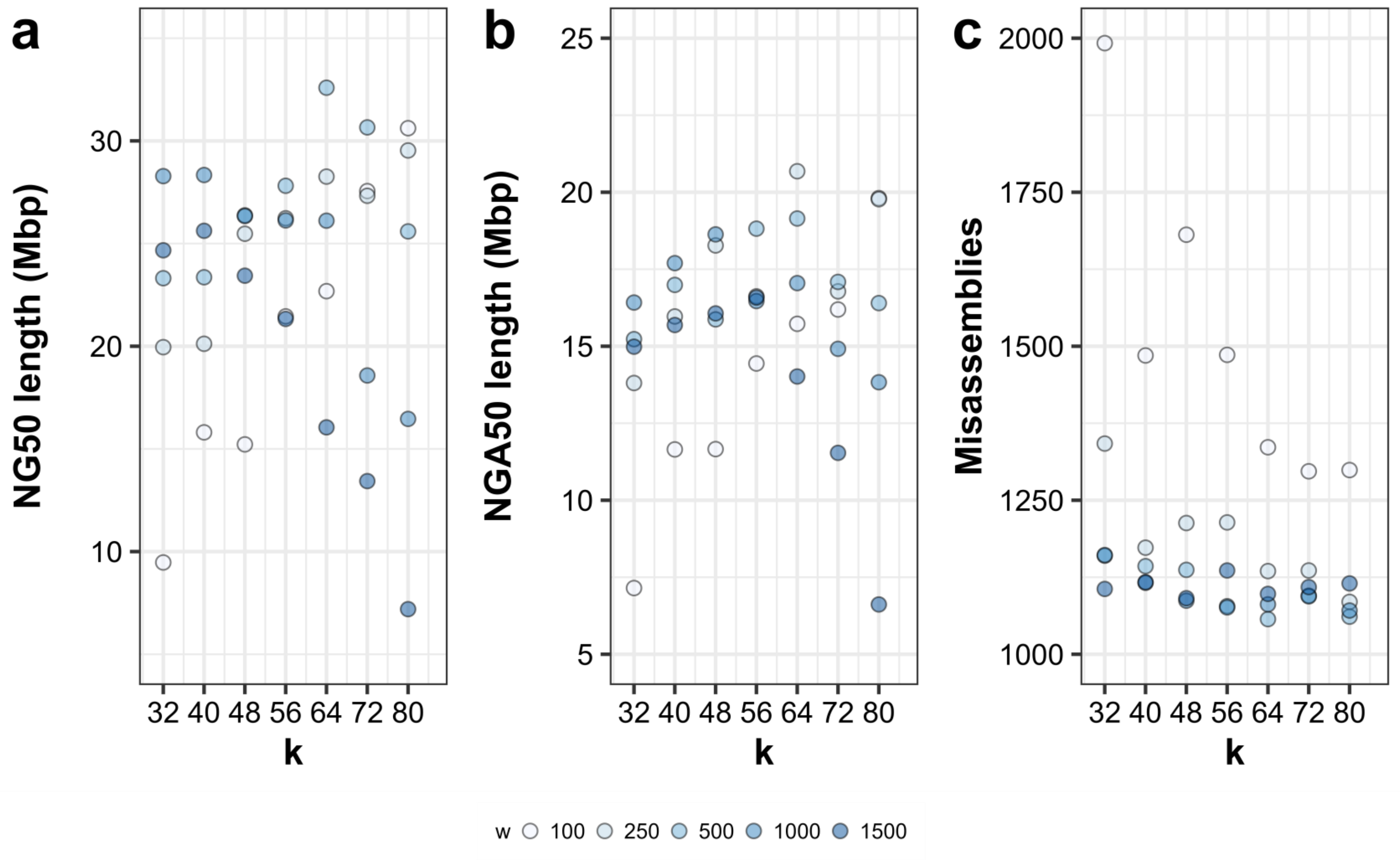

**Supplementary Fig. S10. Contiguity and correctness results of assembling long reads from the human individual HG01243 using GoldRush, sweeping on the GoldRush-Link k and w parameters.**

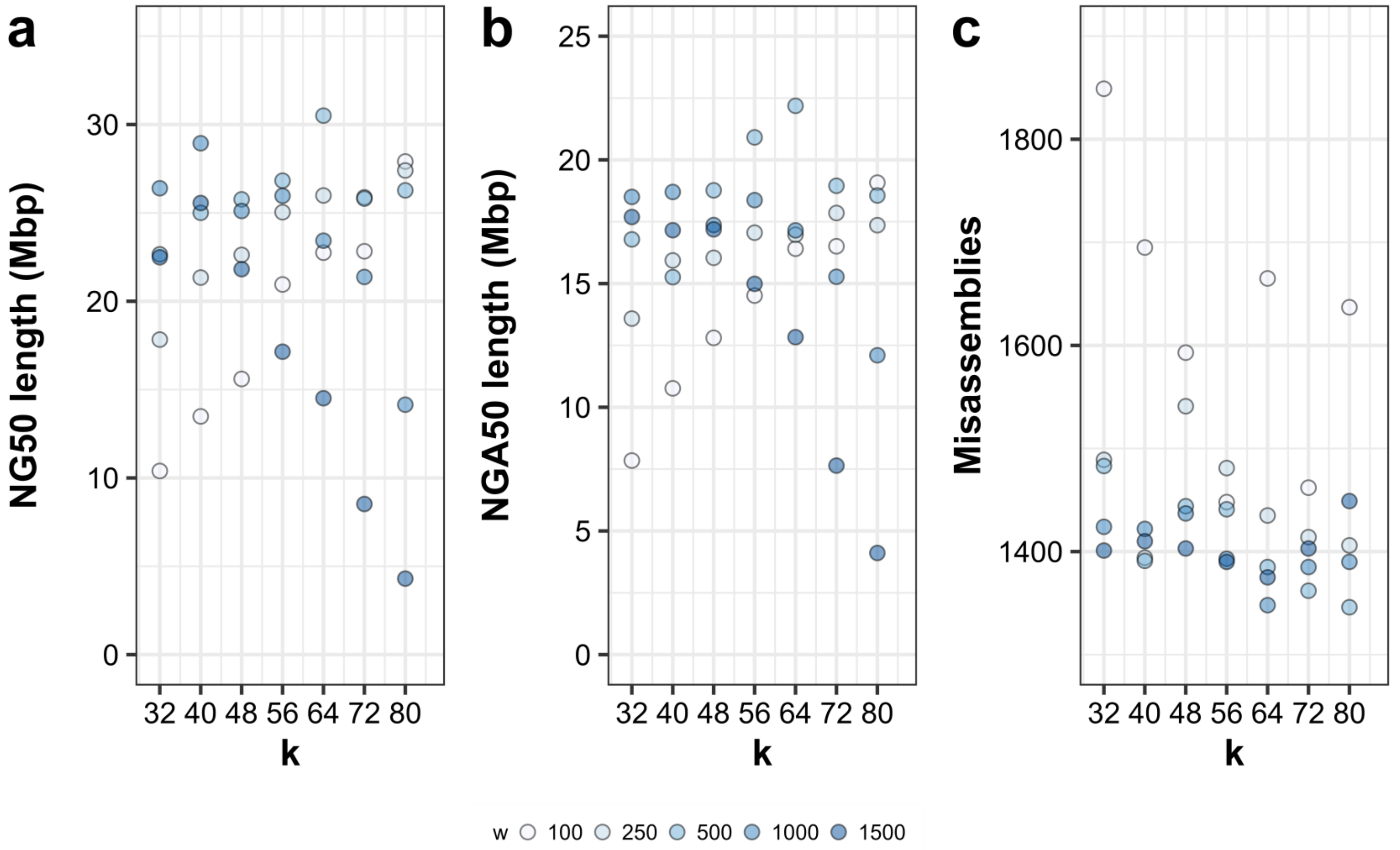

**Supplementary Fig. S11. Contiguity and correctness results of assembling long reads from the human individual HG02055 using GoldRush, sweeping on the GoldRush-Link k and w parameters.**

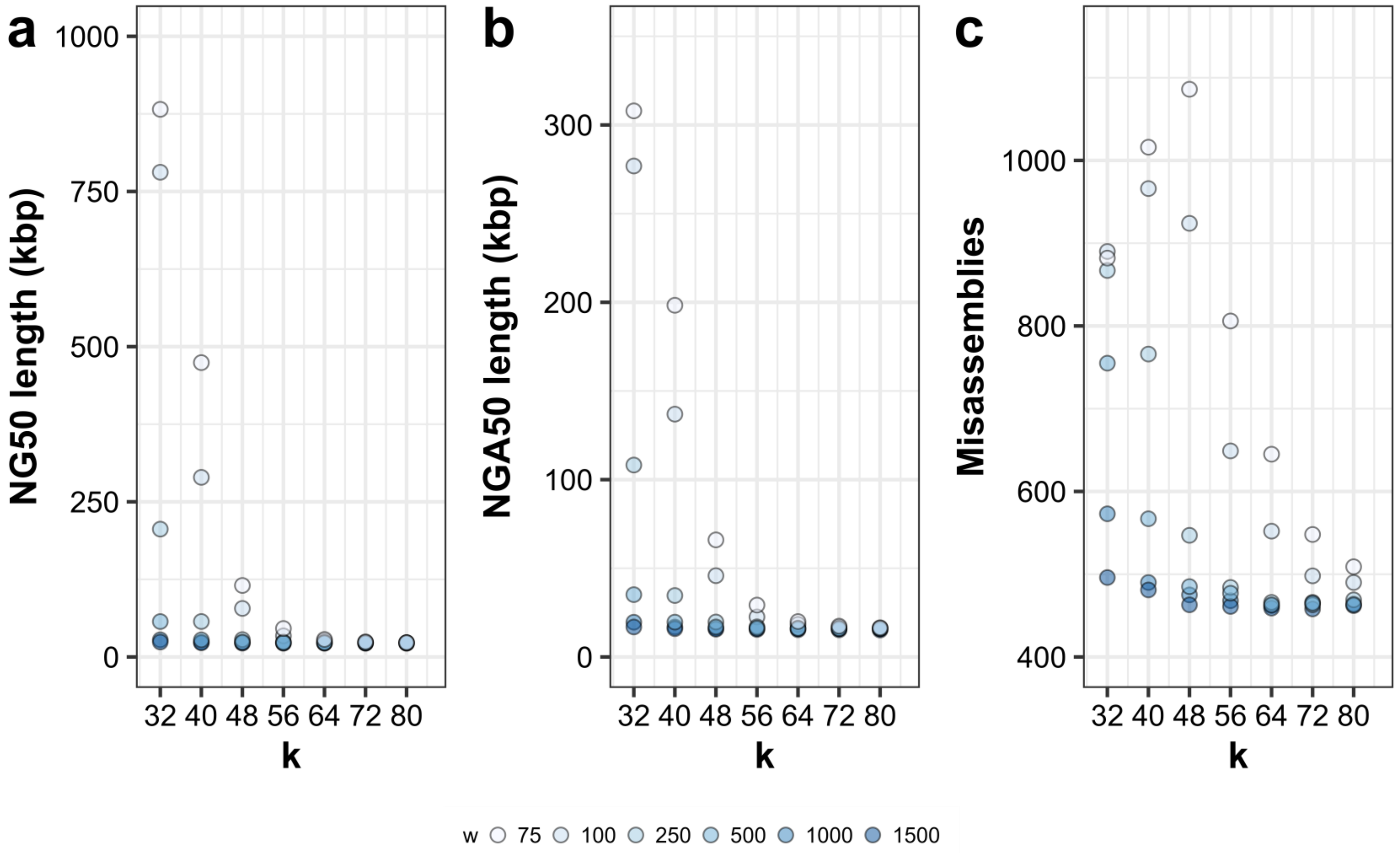

**Supplementary Fig. S12. Contiguity and correctness results of assembling long reads from *O. sativa* using GoldRush, sweeping on the GoldRush-Link k and w parameters.**

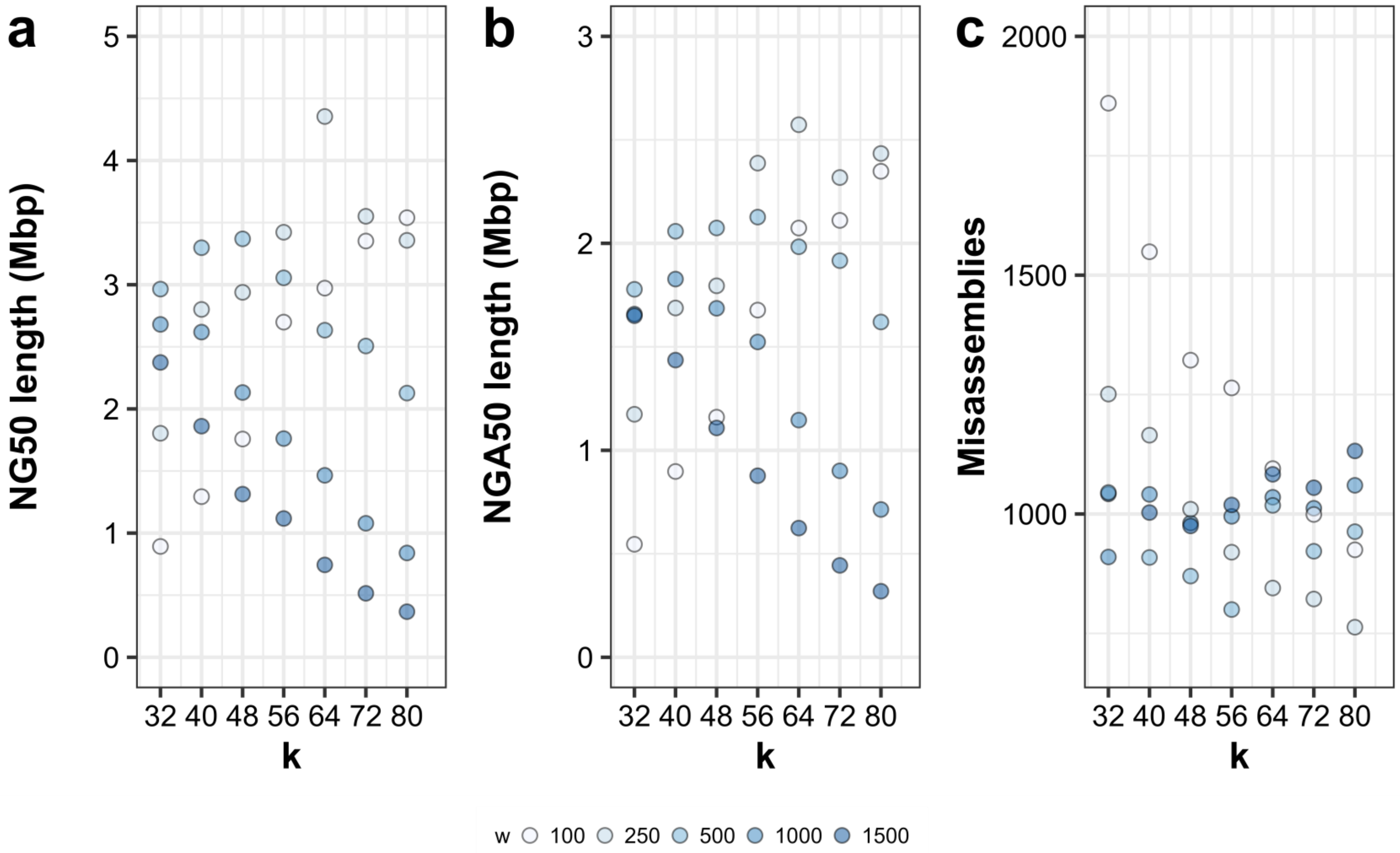

**Supplementary Fig. S13. Contiguity and correctness results of assembling long reads from *S. lycopersicum* using GoldRush, sweeping on the GoldRush-Link k and w parameters.**

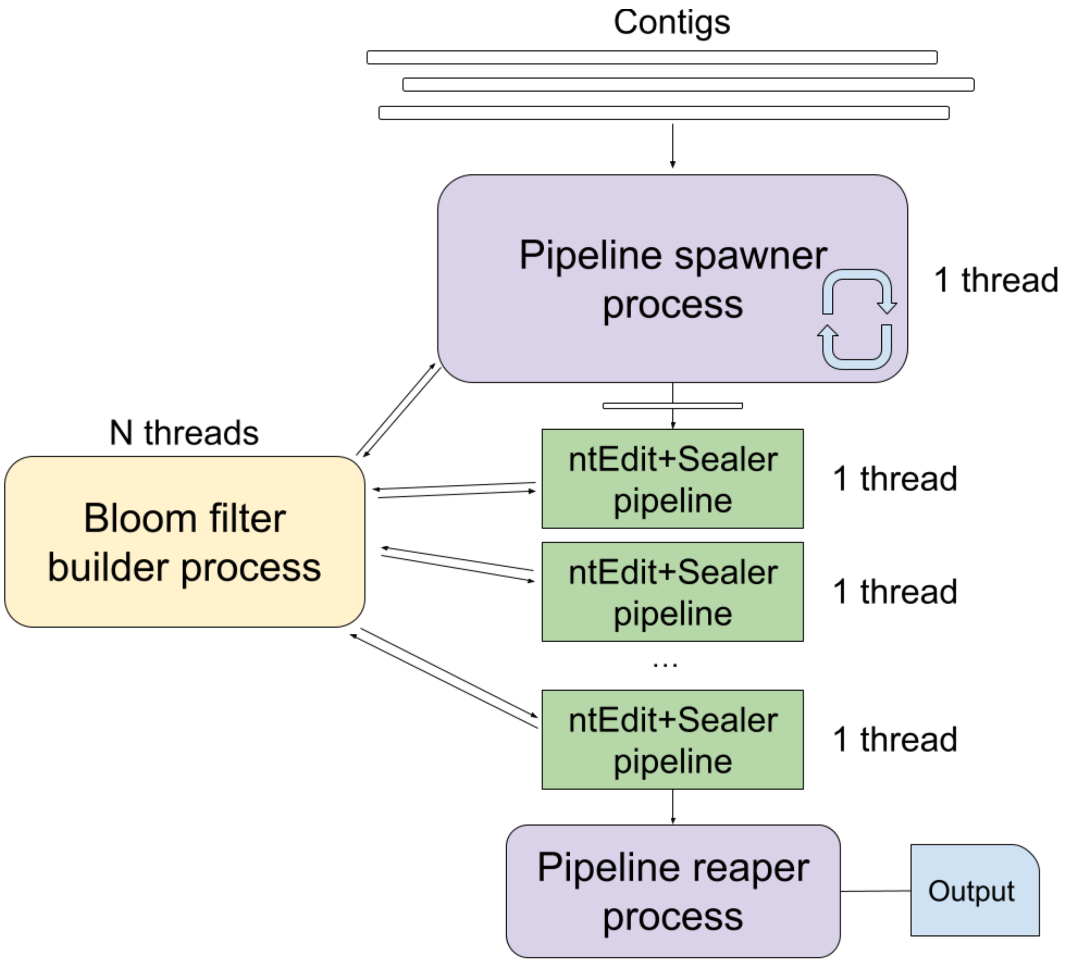

**Supplementary Fig. S14. GoldRush-Edit thread optimization schematic.** GR-Edit coordinates a set of processes in order to maximize concurrency. A pipeline spawner process launches an instance of ntEdit+Sealer per goldtig to polish, up to a specified limit at a time. The Bloom filter builder process continually builds the required Bloom filters for the subsequent polishing pipelines. Finally, a reaper process collects the results of each polishing process in the order they were started and consolidates the polished sequences to a single output fasta file.

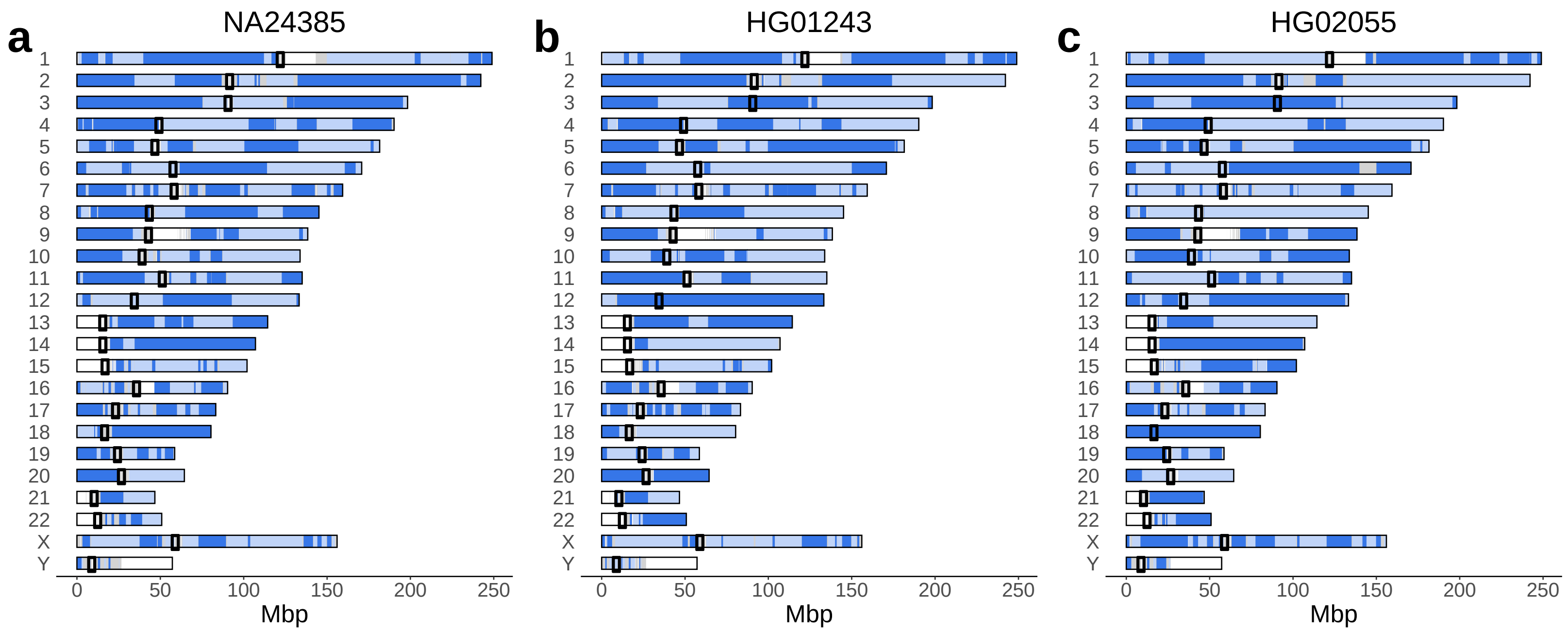

**Supplementary Fig. S15. Ideogram plots showing the contiguity of the three GoldRush human genome assemblies (NA24385, HG01243, and HG02055)**. The ideogram is created by selecting sequences that are least the NG90 length, aligning the sequences to the human reference genome (GRCh38) using minimap2, and plotting the resulting alignment blocks. White regions are Ns in the reference genome, and grey regions have no mapped sequences. The aligned scaffolds are represented with alternating shades of blue. For all three human individuals, GoldRush assembled multiple full chromosome arms. In (a) and (b), GoldRush assembled both chromosomal arms of chromosome 20 and one chromosomal arm of chromosome 18. In (c), GoldRush assembled chromosome 18 in one piece.

**Supplementary Table S1. ONT long read sequencing reads used for genome assembly benchmarks.** All datasets were sequenced using the R9.4.1 chemistry except for the *O. sativa* dataset, which is unknown. The error rate of each dataset was estimated using NanoSim (Yang et al., 2017).

| Species | Individual/Strain | Fold Coverage | N50 Length (bp) | Accession(s)/ Source | Basecaller | Estimated Error rate (%) |
|---|---|---|---|---|---|---|
| ***H. sapiens*** | NA24385 | 67 | 30,348 | s3://ont-open-data/gm24385_2020.11/analysis/r9.4.1/20201026_1644_2-E5-H5_PAG07162_d7f262d5/guppy_v4.0.11_r9.4.1_hac_prom/align_unfiltered/chr1/guppy_v5.0.6_r9.4.1_sup_prom/ | Guppy v5 | 4 |
| ***H. sapiens*** | HG01243 | 63 | 45,310 | https://s3-us-west-2.amazonaws.com/human-pangenomics/index.html?prefix=NHGRI_UCSC_panel/HG02055/nanopore/Guppy_4.2.2/ | Guppy v4 | 9 |
| ***H. sapiens*** | HG02055 | 71 | 47,934 | https://s3-us-west-2.amazonaws.com/human-pangenomics/index.html?prefix=NHGRI_UCSC_panel/HG01243/nanopore/Guppy_4.2.2/ | Guppy v4 | 11 |
| ***O. sativa*** | Japonica group | 62 | 29,349 | SRR10589512-SRR10589711 | - | 20 |
| ***S. lycopersicum*** | MoneybergTMV cultivar | 72 | 23,687 | ERR6668574, ERR7928672 | Guppy v5 | 8 |

"-" denotes unknown.

**Supplementary Table S2. Optimized parameters used for the GoldRush genome assemblies.** Default parameters were used for GoldRush parameters not listed.

| GoldRush Assembly | Optimized Parameters |
|---|---|
| **NA24385** | G=3e9 k_ntLink=72 w_ntLink =500 |
| **HG01243** | G=3e9 k_ntLink=64 w_ntLink =500 |
| **HG02055** | G=3e9 k_ntLink=64 w_ntLink =500 |
| ***O. sativa*** | G=373e6 k_ntLink=32 w_ntLink =75 |
| ***S. lycopersicum*** | G=824e6 k_ntLink=80 |

**Supplementary Table S3. Reference genome builds used for QUAST assembly analysis.**

| Species | Reference genome build | Reference genome accession |
|---|---|---|
| ***H. sapiens*** | GRCh38 | GCA_000001405.15 |
| ***O. sativa*** | IRGSP-1.0 | GCF_001433935.1 |
| ***S. lycopersicum*** | MbTMV | GCA_915070445.1 |

**Supplementary Table S4. Contiguity and correctness statistics of human individual NA24385 genome assemblies generated by GoldRush and the comparator tools.** All statistics were generated using QUAST (Gurevich et al., 2013).

| Assembler | Scaffold NG50 (Mbp) | Scaffold NGA50 (Mbp) | Contig NGA50 (Mbp) | Contig NGA50 (Mbp) | Num. misassemblies | Num. local misassemblies | Genome fraction (%) | Total length (Gbp) | Duplication ratio | Num. mismatches per 100 kbp | Num. indels per 100 kbp | Num. N's per 100 kbp |
|---|---|---|---|---|---|---|---|---|---|---|---|---|
| **GoldRush** | 25.3 | 18.3 | 21.6 | 16.3 | 940 | 1,210 | 94.7 | 2.9 | 1.0 | 257.2 | 227.1 | 53.7 |
| **Flye** | 26.6 | 26.1 | 26.6 | 26.1 | 940 | 1,679 | 96.3 | 2.9 | 1.0 | 140.6 | 54.5 | 0.0 |
| **Redbean** | 8.0 | 7.3 | 8.0 | 7.3 | 4,918 | 2,416 | 92.1 | 2.9 | 1.1 | 255.7 | 154.4 | 0.0 |
| **Shasta** | 29.7 | 24.2 | 29.7 | 24.2 | 1,682 | 2,242 | 95.8 | 2.9 | 1.0 | 150.8 | 57.0 | 0.0 |

**Supplementary Table S5. Contiguity and correctness statistics of human individual HG01243 genome assemblies generated by GoldRush and the comparator tools.** All statistics were generated using QUAST (Gurevich et al., 2013).

| Assembler | Scaffold NG50 (Mbp) | Scaffold NGA50 (Mbp) | Contig NGA50 (Mbp) | Contig NGA50 (Mbp) | Num. misassemblies | Num. local misassemblies | Genome fraction (%) | Total length (Gbp) | Duplication ratio | Num. mismatches per 100 kbp | Num. indels per 100 kbp | Num. N's per 100 kbp |
|---|---|---|---|---|---|---|---|---|---|---|---|---|
| **GoldRush** | 32.6 | 19.1 | 26.7 | 18.0 | 1,057 | 4,010 | 94.5 | 3.0 | 1.1 | 1,387.2 | 1,034.3 | 47.5 |
| **Flye** | 31.3 | 25.6 | 31.3 | 25.6 | 1,062 | 1,473 | 96.2 | 2.9 | 1.0 | 148.7 | 110.6 | 0 |
| **Redbean** | 10.3 | 8.2 | 10.3 | 8.2 | 7,052 | 2,267 | 92.8 | 2.9 | 1.1 | 324.7 | 354.5 | 0 |
| **Shasta** | 35.2 | 30.0 | 35.2 | 30.0 | 3,240 | 3,234 | 96.7 | 2.9 | 1.0 | 195.6 | 100.7 | 0 |

**Supplementary Table S6. Contiguity and correctness statistics of human individual HG02055 genome assemblies generated by GoldRush and the comparator tools.** All statistics were generated using QUAST (Gurevich et al., 2013).

| Assembler | Scaffold NG50 (Mbp) | Scaffold NGA50 (Mbp) | Contig NGA50 (Mbp) | Contig NGA50 (Mbp) | Num. misassemblies | Num. local misassemblies | Genome fraction (%) | Total length (Gbp) | Duplication ratio | Num. mismatches per 100 kbp | Num. indels per 100 kbp | Num. N's per 100 kbp |
|---|---|---|---|---|---|---|---|---|---|---|---|---|
| **GoldRush** | 30.5 | 22.2 | 28.2 | 17.3 | 1,385 | 4,667 | 94.7 | 3.1 | 1.1 | 1,985.0 | 1,415.5 | 57.1 |
| **Flye** | 38.9 | 30.4 | 38.8 | 30.4 | 1,048 | 1,697 | 96.4 | 2.9 | 1.0 | 155.5 | 126.0 | 0.0 |
| **Redbean** | 10.9 | 8.2 | 10.9 | 8.2 | 7,383 | 2,663 | 93.1 | 3.0 | 1.0 | 346.1 | 346.1 | 0.0 |
| **Shasta** | 39.6 | 33.7 | 39.6 | 33.7 | 3,314 | 3,217 | 96.6 | 2.9 | 1.0 | 210.2 | 103.9 | 0.0 |

**Supplementary Table S7. Contiguity and correctness statistics of *O. sativa* genome assemblies generated by GoldRush and the comparator tools.** All statistics were generated using QUAST (Gurevich et al., 2013).

| Assembler | Scaffold NG50 (Mbp) | Scaffold NGA50 (Mbp) | Contig NGA50 (Mbp) | Contig NGA50 (Mbp) | Num. misassemblies | Num. local misassemblies | Genome fraction (%) | Total length (Mbp) | Duplication ratio | Num. mismatches per 100 kbp | Num. indels per 100 kbp | Num. N's per 100 kbp |
|---|---|---|---|---|---|---|---|---|---|---|---|---|
| **GoldRush** | 0.9 | 0.3 | 0.9 | 0.3 | 882 | 16,291 | 78.3 | 451.7 | 1.1 | 8,488.4 | 5,164.9 | 56.7 |
| **Flye** | 3.1 | 1.9 | 2.9 | 1.9 | 679 | 858 | 94.4 | 394.2 | 1.1 | 1,197.6 | 1,201.1 | 0.2 |
| **Redbean** | 5.2 | 2.4 | 5.2 | 2.4 | 483 | 1,663 | 90.0 | 363.2 | 1.0 | 1,344.4 | 2,386.5 | 0.0 |
| **Shasta** | 0.1 | 0.1 | 0.1 | 0.1 | 343 | 3,659 | 71.5 | 284.2 | 1.0 | 2,489.2 | 2,178.5 | 0.0 |

**Supplementary Table S8. Contiguity and correctness statistics of *S. lycopersicum* genome assemblies generated by GoldRush and the comparator tools.** All statistics were generated using QUAST (Gurevich et al., 2013).

| Assembler | Scaffold NG50 (Mbp) | Scaffold NGA50 (Mbp) | Contig NGA50 (Mbp) | Contig NGA50 (Mbp) | Num. misassemblies | Num. local misassemblies | Genome fraction (%) | Total length (Gbp) | Duplication ratio | Num. mismatches per 100 kbp | Num. indels per 100 kbp | Num. N's per 100 kbp |
|---|---|---|---|---|---|---|---|---|---|---|---|---|
| **GoldRush** | 4.4 | 2.6 | 3.9 | 2.5 | 845 | 4,161 | 90.5 | 1.0 | 1.2 | 4,180.9 | 2,052.1 | 96.4 |
| **Flye** | 11.2 | 9.8 | 10.5 | 9.8 | 132 | 244 | 97.1 | 0.8 | 1.0 | 0.2 | 537.5 | 207.4 |
| **Redbean** | 0.3 | 0.2 | 0.3 | 0.2 | 4,895 | 13,726 | 94.8 | 2.2 | 2.7 | 0 | 6,283.2 | 5,633.5 |
| **Shasta** | 26.9 | 21.6 | 26.9 | 21.6 | 82 | 1,062 | 97.0 | 0.8 | 1.0 | 0 | 468.7 | 487.1 |

**Supplementary Table S9. Resource usage of GoldRush and the comparator tools for the genome assembly of the human individual NA24385.** Time and peak memory usage were recorded using the unix *time* command.

| Assembler | Time (h) | Peak memory (GB) |
|---|---|---|
| **GoldRush** | 16.5 | 51.9 |
| **Flye** | 33.7 | 502.4 |
| **Redbean** | 53.3 | 335.1 |
| **Shasta** | 4.4 | 1,003.0 |

**Supplementary Table S10. Resource usage of GoldRush and the comparator tools for the genome assembly of the human individual HG01243.** Time and peak memory usage were recorded using the unix *time* command.

| Assembler | Time (h) | Peak memory (GB) |
|---|---|---|
| **GoldRush** | 20.8 | 53.9 |
| **Flye** | 47.5 | 366.4 |
| **Redbean** | 68.0 | 329.3 |
| **Shasta** | 4.1 | 884.5 |

**Supplementary Table S11. Resource usage of GoldRush and the comparator tools for the genome assembly of the human individual HG02055.** Time and peak memory usage were recorded using the unix *time* command.

| Assembler | Time (h) | Peak memory (GB) |
|---|---|---|
| **GoldRush** | 20.8 | 54.5 |
| **Flye** | 49.2 | 441.0 |
| **Redbean** | 68.1 | 332.9 |
| **Shasta** | 5.0 | 1009.2 |

**Supplementary Table S12. Resource usage of GoldRush and the comparator tools for the genome assembly of the *O. sativa* dataset.** Time and peak memory usage were recorded using the unix *time* command.

| Assembler | Time (h) | Peak memory (GB) |
|---|---|---|
| **GoldRush** | 1.6 | 35.7 |
| **Flye** | 12.1 | 106.3 |
| **Redbean** | 1.4 | 67.8 |
| **Shasta** | 0.5 | 98.1 |

**Supplementary Table S13. Resource usage of GoldRush and the comparator tools for the genome assembly of the *S. lycopersicum* dataset.** Time and peak memory usage were recorded using the unix *time* command.

| Assembler | Time (h) | Peak memory (GB) |
|---|---|---|
| **GoldRush** | 7.4 | 45.3 |
| **Flye** | 24.6 | 255.0 |
| **Redbean** | 14.1 | 160.6 |
| **Shasta** | 2.3 | 262.5 |

**Supplementary Table S14. Contiguity and correctness statistics of human individual NA24385 genome assembly generated by GoldRush at different stages.** All statistics were generated using QUAST (Gurevich et al., 2013).

| Stage | Scaffold NG50 (Mbp) | Scaffold NGA50 (Mbp) | Contig NGA50 (Mbp) | Contig NGA50 (Mbp) | Num. misassemblies | Num. local misassemblies | Genome fraction (%) | Total length (Gbp) | Duplication ratio | Num. mismatches per 100 kbp | Num. indels per 100 kbp | Num. N's per 100 kbp |
|---|---|---|---|---|---|---|---|---|---|---|---|---|
| **GR-Path** | 0.02 | 0.02 | 0.02 | 0.02 | 2,948 | 1,994 | 92.7 | 2.9 | 1.1 | 1,463.7 | 1,327.2 | 0.0 |
| **GR-Edit** | 0.02 | 0.02 | 0.02 | 0.02 | 2,990 | 1,777 | 92.7 | 2.9 | 1.1 | 228.5 | 197.2 | 23.4 |
| **Tigmint-Long** | 0.02 | 0.02 | 0.02 | 0.02 | 1,863 | 900 | 92.6 | 2.9 | 1.1 | 224.1 | 196.1 | 23.2 |
| **GR-Link** | 25.3 | 18.3 | 21.6 | 16.3 | 940 | 1,210 | 94.8 | 2.8 | 1.0 | 257.2 | 227.1 | 53.7 |

**Supplementary Table S15. BUSCO statistics for assemblies of human individual NA24385 generated by GoldRush and the comparator tools.** BUSCO was run using the primates_odb10 lineage (Simão et al., 2015).

| Assembler | Complete BUSCOs | Complete and single-copy BUSCOs | Complete and duplicated BUSCOs | Fragmented BUSCOs | Missing BUSCOs | Total BUSCO groups searched |
|---|---|---|---|---|---|---|
| **GoldRush** | 12,272 (89.1%) | 12,059 | 213 | 511 | 997 | 13,780 |
| **Flye** | 12,988 (94.3%) | 12,769 | 219 | 260 | 532 | 13,780 |
| **Redbean** | 12,193 (88.5%) | 11,967 | 226 | 412 | 1175 | 13,780 |
| **Shasta** | 12,920 (93.8%) | 12,703 | 217 | 199 | 661 | 13,780 |

**Supplementary Table S16. BUSCO statistics for assemblies of human individual HG01243 generated by GoldRush and the comparator tools.** BUSCO was run using the primates_odb10 lineage (Simão et al., 2015).

| Assembler | Complete BUSCOs | Complete and single-copy BUSCO | Complete and duplicated BUSCO | Fragmented BUSCO | Missing BUSCOs | Total BUSCO groups searched |
|---|---|---|---|---|---|---|
| **GoldRush** | 8,413 (61.1%) | 8,295 | 118 | 597 | 4,770 | 13,780 |
| **Flye** | 12,344 (89.6%) | 12,130 | 214 | 526 | 910 | 13,780 |
| **Redbean** | 10,964 (79.6%) | 10,789 | 175 | 725 | 2,091 | 13,780 |
| **Shasta** | 12,485 (90.6%) | 12,279 | 206 | 464 | 831 | 13,780 |

**Supplementary Table S17. BUSCO statistics for assemblies of human individual HG02055 generated by GoldRush and the comparator tools.** BUSCO was run using the primates_odb10 lineage (Simão et al., 2015).

| Assembler | Complete BUSCOs | Complete and single-copy BUSCOs | Complete and duplicated BUSCOs | Fragmented BUSCOs | Missing BUSCOs | Total BUSCO groups searched |
|---|---|---|---|---|---|---|
| **GoldRush** | 7,108 (51.6%) | 7,016 | 92 | 569 | 6,103 | 13,780 |
| **Flye** | 12,215 (88.6%) | 11,985 | 230 | 559 | 1,006 | 13,780 |
| **Redbean** | 10,763 (78.1%) | 10,592 | 171 | 669 | 2,348 | 13,780 |
| **Shasta** | 12,471 (90.5%) | 12,261 | 210 | 451 | 858 | 13,780 |

**Supplementary Table S18. Contiguity and correctness statistics of the GoldRush genome assembly of the human individual NA24385 using Racon instead of GoldRush-Edit at different stages.** All statistics were generated using QUAST (Gurevich et al., 2013).

| Stage | Scaffold NG50 (Mbp) | Scaffold NGA50 (Mbp) | Contig NGA50 (Mbp) | Contig NGA50 (Mbp) | Num. misassemblies | Num. local misassemblies | Genome fraction (%) | Total length (Gbp) | Duplication ratio | Num. mismatches per 100 kbp | Num. indels per 100 kbp | Num. N's per 100 kbp |
|---|---|---|---|---|---|---|---|---|---|---|---|---|
| **GR-Path** | 0.02 | 0.02 | 0.02 | 0.02 | 2,948 | 1,994 | 92.7 | 2.9 | 1.1 | 1,463.7 | 1,327.2 | 0.0 |
| **Racon** | 0.02 | 0.02 | 0.02 | 0.02 | 2,750 | 479 | 92.8 | 2.9 | 1.1 | 157.0 | 106.4 | 0.0 |
| **Tigmint-Long** | 0.02 | 0.02 | 0.02 | 0.02 | 1,665 | 410 | 92.7 | 2.9 | 1.1 | 155.8 | 105.9 | 0.0 |
| **GR-Link** | 24.4 | 14.9 | 24.0 | 14.4 | 945 | 589 | 94.6 | 2.9 | 1.0 | 185.0 | 134.2 | 33.8 |

**Supplementary Table S19. BUSCO statistics for assemblies of human individual NA24385 generated by GoldRush using Racon instead of GoldRush-Edit.** BUSCO was run using the primates_odb10 lineage (Simão et al., 2015).

| Complete BUSCOs | Complete and single-copy BUSCOs | Complete and duplicated BUSCOs | Fragmented BUSCOs | Missing BUSCOs | Total BUSCO groups searched |
|---|---|---|---|---|---|
| 12,752 (92.5%) | 12,533 | 219 | 296 | 732 | 13,780 |

**Supplementary Table S20. Resource usage breakdown of each GoldRush stage for the genome assembly of human individual NA24385.** Time and peak memory usage were recorded using the unix *time* command.

| Stage | Time (h) | Peak memory (GB) |
|---|---|---|
| **GR-Path** | 3.2 | 51.9 |
| **GR-Edit** | 8.3 | 11.0 |
| **Tigmint-Long** | 1.0 | 11.4 |
| **GR-Link** | 4.1 | 42.9 |
| **TOTAL GoldRush** | 16.6 | 51.9 |

**Supplementary Table S21. Resource usage breakdown of each GoldRush stage for the genome assembly of the human individual HG01243.** Time and peak memory usage were recorded using the unix *time* command.

| Stage | Time (h) | Peak memory (GB) |
|---|---|---|
| **GR-Path** | 4.3 | 53.9 |
| **GR-Edit** | 12.0 | 12.3 |
| **Tigmint-Long** | 0.8 | 11.1 |
| **GR-Link** | 3.7 | 26.3 |
| **TOTAL GoldRush** | 20.8 | 53.9 |

**Supplementary Table S22. Resource usage breakdown of each GoldRush stage for the genome assembly of the human individual HG02055.** Time and peak memory usage were recorded using the unix *time* command.

| Stage | Time (h) | Peak memory (GB) |
|---|---|---|
| **GR-Path** | 3.6 | 54.5 |
| **GR-Edit** | 12.5 | 13.3 |
| **Tigmint-Long** | 0.9 | 11.7 |
| **GR-Link** | 4.0 | 25.8 |
| **TOTAL GoldRush** | 21.0 | 54.5 |

**Supplementary Method S1. Improving the accuracy of the best hits for tiles.**

After the preliminary hits are established, the tiles are sequentially re-analyzed. The ID of the current tile is compared to the ID of the previous tile, and if the IDs do not match, the ID of the previous tile is queried against the current tile's ID-to-counts table. If the ID of the previous tile is found in the current tile's ID-to-counts table, the current tile's associated ID will be changed to that ID and the assignment will be adjusted depending on whether the newly changed ID has a count greater than the threshold (Supplementary Fig. S4a). Next, all the unassigned tiles in the read will be targeted. These tiles will be compared against their two adjacent tiles to see if the adjacent tiles share the same ID or have an ID that is 1 greater or smaller. If the adjacent tiles have the expected associated ID and these tiles are assigned, the current unassigned tile will be assigned (Supplementary Fig. S4b). After single unassigned tiles flanked by two assigned tiles are resolved, stretches of unassigned tiles flank by two assigned tiles will be targeted. If the assigned tiles flanking the unassigned region have the same ID or an ID that is 1 greater or smaller than the other, all the unassigned tiles that are flanked by the two tiles will be assigned and given the associated ID of either flank (Supplementary Fig. S4c). Finally, isolated assigned tiles are selected, and their assignment are changed to unassigned (Supplementary Fig. S4d).
